# Thymic mesenchymal niche cells drive T cell immune regeneration

**DOI:** 10.1101/2022.10.12.511184

**Authors:** Karin Gustafsson, Sergey Isaev, Kameron A. Kooshesh, Ninib Baryawno, Konstantinos D. Kokkaliaris, Nicolas Severe, Ting Zhao, Elizabeth W. Scadden, Joel A. Spencer, Christian Burns, Kumaran Akilan, Nikolaos Barkas, Hayalneh Gessessew, Charles P. Lin, Peter V. Kharchenko, David T. Scadden

**Author notes:** Corresponding authors: David T. Scadden, Karin Gustafsson. Altos Labs, San Diego, CA 92121.

## Abstract

Thymic atrophy and the progressive immune decline that accompanies it is a major health problem, chronically with age and acutely with immune injury. No solution has been defined. Here we demonstrate that one of the three mesenchymal cell subsets identified by single-cell analysis of human and mouse thymic stroma is a critical niche component for T lymphopoiesis. The Postn+ subset is located perivascularly in the cortical-medullary junction, medulla and subcapsular regions. Cell depletion demonstrated that it recruits T competent cells to the thymus and initiates T lymphopoiesis *in vivo*. This subset distinctively expresses the chemokine *Ccl19* necessary for niche functions. It markedly declines with age and in the acute setting of hematopoietic stem cell transplant conditioning. When isolated and adoptively transferred, these cells durably engrafted the atrophic thymus, recruited early T progenitors, increased T cell neogenesis, expanded TCR complexity and enhanced T cell response to vaccination. These data define a thymus lymphopoietic niche cell type that may be manipulated therapeutically to regenerate T lymphopoiesis.

## Introduction

Maintenance of a diverse T cell population is essential for immune homeostasis. Inherited and acquired T cell deficiencies cause highly morbid diseases. Notably, the immune senescence of aging is associated with increased infectious, autoimmune disorders and oncologic disease ^1-3^. Acutely, failure to adequately regenerate T cell immunity after hematopoietic stem cell transplantation (HSCT) is a major cause of morbidity and mortality ^4^. HSCT patients that achieve rapid T cell recovery are less likely to relapse, succumb to lethal infections and suffer severe graft-versus-host disease ^5-7^. Preserving or reestablishing T cell production is therefore an issue of medical importance.

T cell development is dependent on influx of bone marrow progenitors as well as on the stromal cell compartment of the thymus ^8,9^. Thymic epithelial cells (TECs) have been shown to sustain T cell production ^10,11^ and partake in antigenic selection ^12,13^. Endothelial cells secrete cytokines supporting hematopoietic and stroma constituents ^14,15^. Most previous work on improving thymus stromal cell function has focused on enhancing TEC recovery ^15,16^ or endothelial cell function ^15^. Yet, thymic mesenchymal cells (ThyMC) are known to be essential in embryonic development providing signals for the recruitment and differentiation of epithelial progenitors. The thymic primordium in the third pharyngeal pouch does not form without them ^17-21^. In the adult, they express lymphopoietic factors like Ccl19 ^22,23^, dendritic cell chemoattractant Cxcl14 ^24^; they maintain medullary thymic epithelial cells ^25^ and are critical for central tolerance ^26^. We characterized thymic mesenchymal functional subsets in humans and mice across different stress-states. We defined three novel ThyMCs by single cell analysis, one of which is distinctly T cell supportive. The T cell supportive ThyMCs express *Ccl19* and are lost upon aging and HSCT. We further demonstrate that transfer of T cell supportive ThyMCs or their gene product rejuvenates thymic tissue resulting in regeneration of T lymphopoiesis, TCR diversity and neoantigen response in aged or acutely injured animals.

## Results

### Single-cell sequencing of human and mouse thymus identifies mesenchymal cell subsets with distinct T cell supportive signatures

To define thymic stromal components in homeostasis, non-hematopoietic, non-erythroid cells from human and murine thymi were isolated and single-cell RNA sequencing (scRNA-seq) performed (Fig 1A, Supplemental fig 1A-B, Supplementary table 1). Following quality control filtering (Supplemental fig 1C) and contaminating hematopoietic cell removal (Supplemental fig 1D-G), 8628 human and 5451 murine stromal cells remained (Fig 1B-C). As has been described by others ^27,28^, both human and mouse thymus contains nine distinct stromal cell subsets (Fig 1B-C, Supplemental fig 2A-D). The human stromal cell compartment consist of endothelial cells (EC; arteriolar, post-capillary and lymphatic), thymic mesenchymal cells (ThyMCs), thymic epithelial cells (TECs; cortical, medullary and myo/neuro), and perivascular cells (pericytes; PC and vascular smooth muscle cells; vSMC) (Fig 1B, Supplemental fig 2A-B). Similarly, the murine stroma contains ThyMCs and two perivascular subsets (Fig 1C, Supplemental fig 2C-D). Yet, some reduction in complexity was observed. The murine samples show only one EC cluster, and TECs are subdivided into just two populations, a mixed population of cortical TEC and immature medullary TEC (cTEC/mTECimm) and a subset of apparent mature medullary TECs (mTECm) (Fig 1C, Supplemental fig 2C-D). Notably, the recently described thymic Tuft cells ^27^ form a distinct population in mouse samples, but no clearly defined cluster was seen in human thymus, in line with another scRNA-seq study ^28^. There were also two small populations only found in mice. A subset expressing the mesothelial marker *Lrrn4* (Fig 1B-C, Supplemental fig 2C-D) ^29^ as well as a small cluster of cells defined by neural crest genes (*Dct, So×10*) ^30^ (Fig 1B-C, Supplemental fig 2C and E). Neural crest derived cells (NC) have previously been demonstrated to give rise to thymic pericytes suggesting that these cells may represent a tissue resident progenitor population ^31,32^.

**Figure 1.**
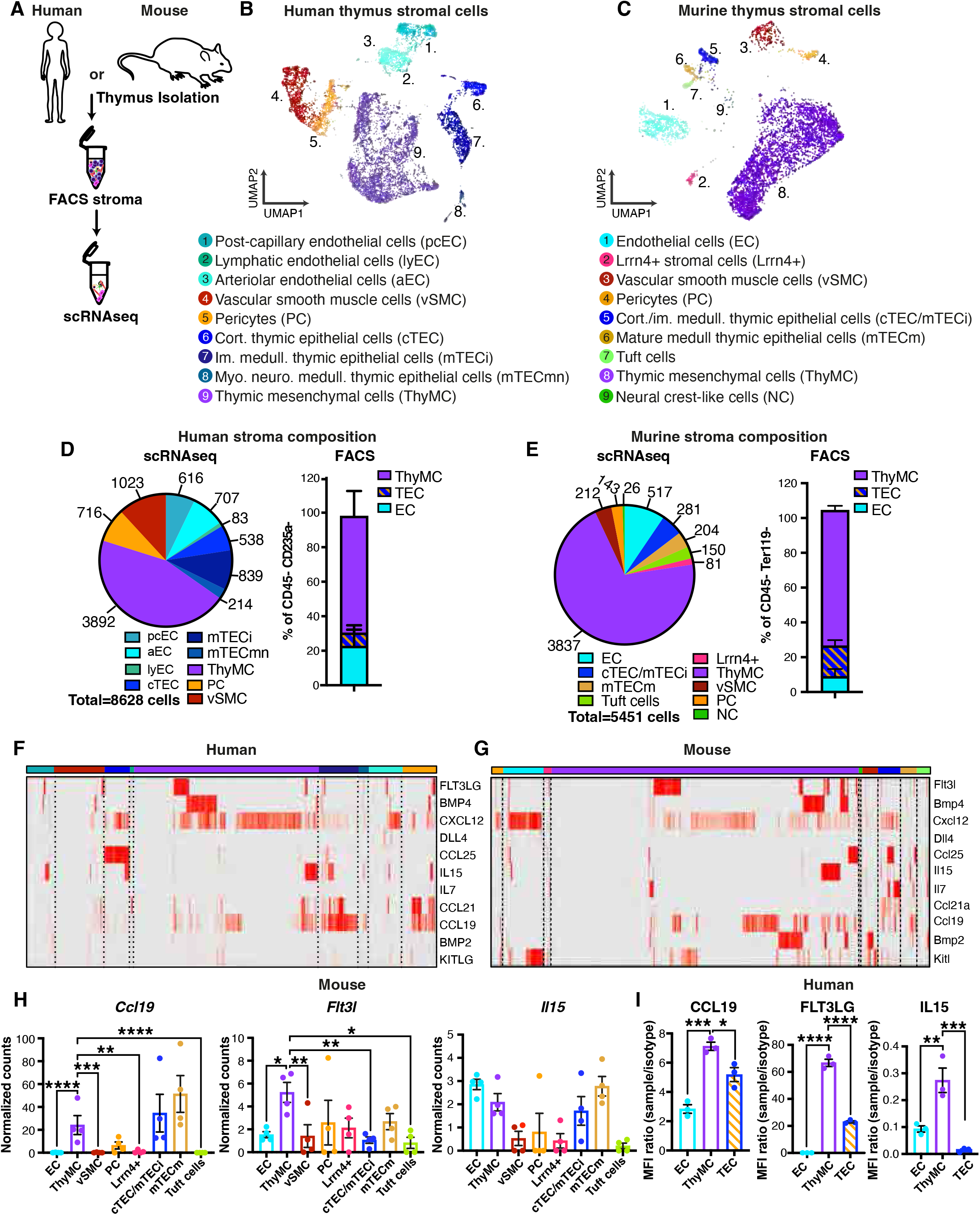
Single-cell sequencing of human and mouse thymus identifies mesenchymal cell subsets with distinct T cell supportive signatures (A) Schematic illustration of human and mouse sample collection. (B) Joint UMAP embedding of human thymus stromal cell populations (*CD3E-, CD4-, CD8B-, PTPRC-* cells) with detailed annotation. (n=3) Three independent experiments, total number of stromal cells is 8628. (C) Joint UMAP embedding of murine thymus stromal cell populations (*Cd3e-, Cd4-, CD8a-, Ptprc-* cells) with detailed annotation. (n=4) Four independent experiments, total number of stromal cells is 5451. (D) Human thymus stromal cell composition as determined by scRNA-seq (left) and flow cytometric analysis (right). (n=3) Three independent experiments. (E) Murine thymus stromal cell composition as determined by scRNA-seq (left) and flow cytometric analysis (right). (n=4) Four independent experiments. (F) Heatmap showing expression of selected lymphopoietic factors (rows) from different human thymus stromal cell populations (columns). (n=3) Three independent experiments. (G) Heatmap showing expression of selected lymphopoietic factors (rows) from different murine stromal cell populations (columns). (n=4) NCs were excluded from this analysis due to the populations small size (6.5 cells per sample). Four independent experiments. (H) Expression of Il15, Ccl19 and Flt3l across different murine stromal cell subsets. Statistical significance was assessed using DESeq2 with Benjamini-Hochberg FDR control of these comparisons. (n=4) Four independent experiments. (I) Bar graphs showing the results of flow cytometric quantification of IL15, CCL19 and FLT3LG in human thymus stromal cell subsets. Data is normalized against the respective isotype control of each cytokine specific antibody. Statistical significance was assessed using one-way ANOVA followed by Dunnet’s post-hoc analysis. (n=3)

The largest stromal cell fraction in both human and mouse was *PRRX1+*^33^, extracellular matrix expressing ThyMCs as confirmed by flow cytometry (Fig 1D-E, Supplemental fig 2A- D). Interestingly, both human and murine thymic ThyMCs expressed high levels of several lymphoid development regulators including *CXCL12* ^34^, *FLT3LG* ^35^, *KITLG* ^14^, *CCL19* ^36^, *IL15* ^37^ and *BMP4* ^15^ (Fig 1F and G). Indeed, murine ThyMCs were among the highest expressors of *IL-15, Flt3l*, and *Ccl19* (Fig 1H). Protein levels of IL15, CCL19 and FLT3LG were confirmed in human thymus samples and exceeded levels in endothelium and epithelium (Fig 1I). These data suggest a functional role for ThyMCs in the lymphopoietic niche.

### Postn+ thymic ThyMCs preferentially express T cell regulators

Three distinct ThyMC subpopulations were evident in both human and murine thymus (Fig 2A, Supplemental fig 3A). Both species were found to have CD248+ ^38^ and Postn+ ThyMC populations (Fig 2A, Supplemental fig 3A-B). The third ThyMC subset was found to be characterized by *COL15A1* expression in humans and *Penk* expression in mouse (Fig 2A, Supplemental fig 3A-B). The correspondence of transcriptional states between human and mouse ThyMCs was confirmed by joint clustering of the ThyMC compartments (Fig 2B).

**Figure 2.**
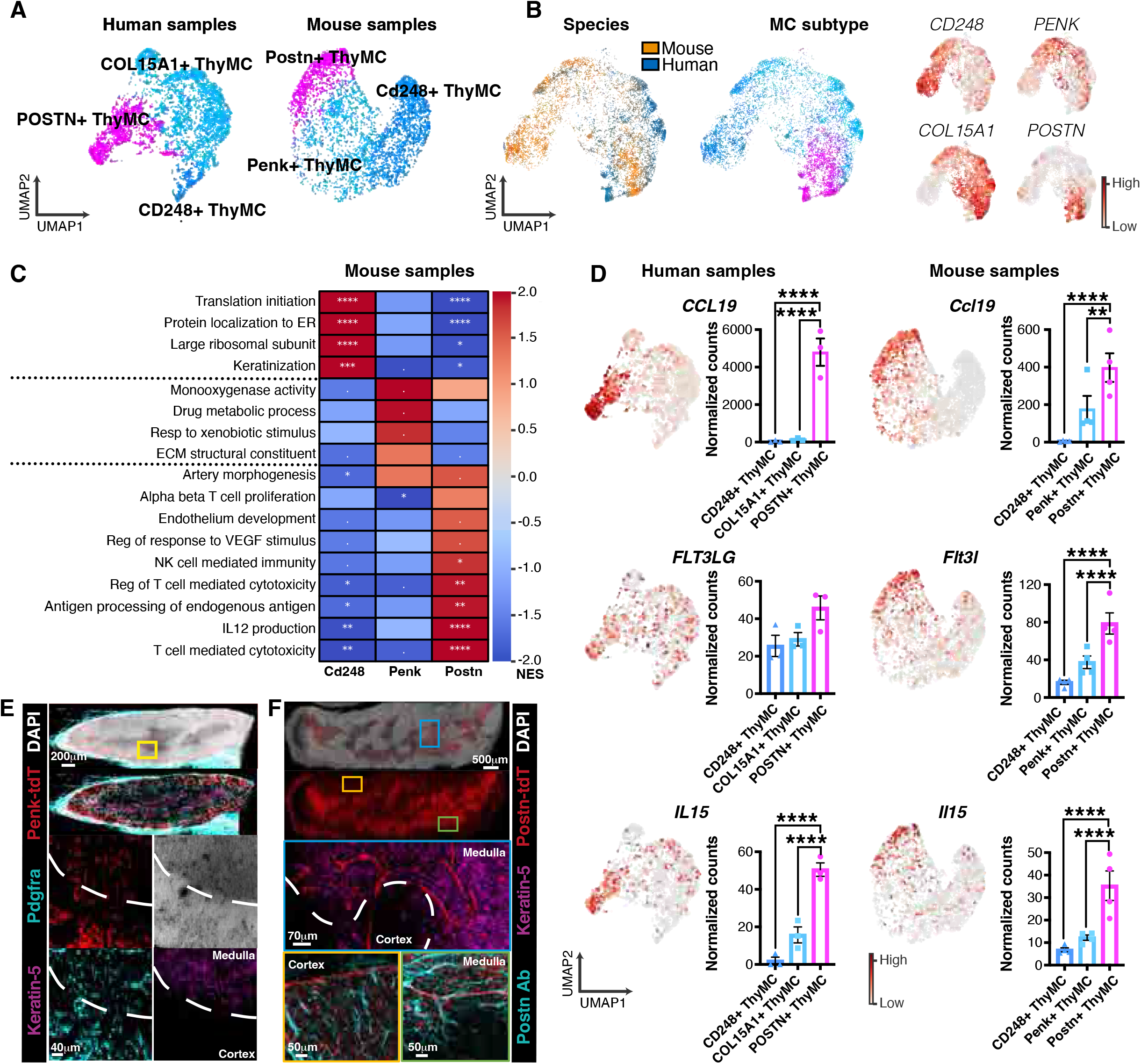
Postn+ thymic ThyMCs preferentially express T cell regulators (A) Detailed analysis and annotation of mesenchymal cell (ThyMC) populations in human (left) and murine (right) samples shown as mesenchymal cell-specific UMAP embedding. (human n = 3 with total number of ThyMCs = 3892, murine n = 4 with total number of ThyMCs = 3837). Cell clusters were identified using unsupervised clustering algorithms. (B) UMAP representation of conos joint graph of human and mouse ThyMCs colored by origin (left). Overlays of individual population markers (right) and ThyMC subtypes (center) indicate transcriptionally similar populations between the two species. (C) Heatmap displaying gene set enrichment analysis (GESA) of differentially expressed genes in murine ThyMC subtypes. Significance level marked as dot corresponds to FDR between 0.05 and 0.25 following the official GSEA User Guide (n=4). (D) Expression of Il15, Ccl19 and Flt3l across the different ThyMC subsets in human and murine samples. Statistical significance was assessed using the DESeq2 with Benjamini- Hochberg FDR control of these comparisons. (human n= 3, murine n= 4). (E) Volumetric confocal microscopy images from a Penk-Cre-tdTomato thymus stained with DAPI (white; cell nuclei), Keratin-5 (purple; mTECs), Pdgfra (cyan; thymic ThyMCs) and tdTomato (red; Penk+ ThyMCs). Dashed line represents the corticomedullary junction. (F) Volumetric confocal microscopy images from a Postn-CreER-tdTomato thymus stained with DAPI (white; cell nuclei), Keratin-5 (purple; mTECs), Periostin (cyan; native protein) and tdTomato (red; Postn+ ThyMCs). Dashed line represents the corticomedullary junction.

GSEA suggested distinct functions among the ThyMC subtypes. CD248+ ThyMCs were enriched for protein translation and extracellular matrix production (ECM) (Fig 2C)) and expressed matrix components, *Fn1* and *Ogn*, (Supplemental fig 3A) consistent with fibroblasts. Penk+ ThyMCs on the other hand, had diverse functional terms, indicating high metabolic activity as well as ECM secretion (Fig 2C). Lastly, Postn+ ThyMCs expressed angiogenesis and abundant T and NK cell associated terms (Fig 2C, Supplemental fig 3F). Additionally, both human and murine Postn+ ThyMCs expressed lymphopoietic cytokines *CCL19* (*Ccl19*), *FLT3LG* (*Flt3l*) and *IL15*, (*Il15*) at significantly higher levels than most other ThyMC subpopulations (Fig 2D) as well as other stromal cells (Supplemental fig 3G). This suggests that Postn ThyMCs may be responsible for the majority of interactions between the mesenchymal compartment and the developing lymphocytes.

Alignment with publicly available datasets ^27,28^ demonstrated presence of three transcriptionally similar human and murine ThyMC subsets (Supplemental fig 3C-D). There were some differences in marker gene expression in the human data, which might be explained by tissue source, the samples produced by Park et. al ^28^ were from embryonic thymi whereas our samples were strictly postnatal. Yet *Ccl19* expressing ThyMCs were readily identified confirming that T cell supportive ThyMCs can be robustly identified in independent samples (Supplemental fig 3C-D).

Seeking markers to prospectively isolate ThyMCs subpopulations, *CD99l2* and *Itgb5* were abundant surface proteins across all three ThyMC subsets, less expressed on other cell types (Supplemental fig 4A), and validated by flow cytometry (Supplemental fig 4B-C) and colony forming ability (Supplemental fig 4D-F). Both Penk+ ThyMCs and Postn+ ThyMCs express *Pdgfra* and lack *Cd248* (Supplemental fig 4A and G) thus CD99l2+Itgb5+Pdgfra+ CD248- cells exclude the fibroblastic CD248+ ThyMCs while enriching for the T cell supportive Postn+ ThyMCs (Supplemental fig 4H). To refine the separation of murine thymic ThyMC subsets we intercrossed *Penk-Cre* or *Postn-CreER* strains with the *Rosa26-LSL-tdTomato* (*tdT)* reporter. Flow cytometric assessment (Supplemental fig 5A-B, E-F) and qPCR analysis of reporter labeled cells (Supplemental fig 5D and G) validated the fidelity of the reporters. A subpopulation of tdTomato+ CD45+ cells were observed in *Penk-Cre-tdT* mice (Supplemental fig 5B). However, these cells were characterized by myeloid cell markers (Supplemental fig 5B) and subsequent intracellular staining for Cre revealed that the cells did not contain Cre protein (Supplemental fig 5C). The tdTomato labeling of this population is thus likely due to phagocytosis of fluorescent material. This suggests that both *Penk-Cre-tdT* and *Postn-CreER-tdT* mice can be used to distinguish T cell supportive Postn+ ThyMCs.

Volumetric confocal tissue imaging of *Penk-Cre-tdT* and *Postn-CreER-tdT* thymi further demonstrated that Penk+ and Postn+ ThyMCs represent anatomically distinct ThyMC populations (Fig 2E-F). Penk+ Pdgfra+ cells exhibited fibroblastic morphology and were distributed interstitially in both the cortex and the medulla (Fig 2E). On the other hand, Postn+ cells were mostly perivascular, and although enriched in the medulla, the cells are also present around cortical capillaries. (Fig 2F). Co-staining with Periostin antibody revealed anatomical proximity between the Periostin+ extracellular matrix network and the Postn-tdT ThyMCs (Fig 2F) as previously reported for other ECM proteins ^39^, validating the faithfulness of the reporter. The presence of Postn+ ThyMCs in both cortex and medulla further suggests that these cells are part of the early lymphoid progenitor niche.

Application of previously used ThyMC markers revealed one or at most two, but never three subsets of cells (Supplemental fig 5H) ^25,26^. Further validation in *Penk-Cre-tdT* mice confirmed this (Supplemental fig 5I-L). Flow cytometric analysis of Pdpn and Dpp4 has recently been used to identify cortical and medullary fibroblasts ^26^. Although both CD248+ and Penk+ ThyMC appear enriched in the DPP4+ Pdpn+ capsular fibroblast (cFib) phenotype, our imaging of the *Penk-Cre-tdT* reporter suggests that these cells are abundant in both cortex and medulla (Fig 2E, Supplemental fig 5K-L). Additionally, Postn+ ThyMC do not show any preferential expression of Pdpn and Dpp4, suggesting that identification of T cell supportive ThyMC will not be possible using this specific set of markers (Supplemental fig 5K-L).

### Postn+ ThyMCs are depleted with acute transplant conditioning and chronic aging

To define stromal cell participation in thymic dysfunction and regeneration, CD45- Ter119- thymic cells were sorted from radiation conditioned HSCT recipients (3 days post-transplantation, acute thymic injury), IL7 receptor knockout mice (IL7RKO) (chronic lymphoid progenitor deficiency) and 2-year old mice (chronic age-associated functional decline) (Fig 3A). Only *Cd3e-, Cd4-, Cd8a-*, and *Ptprc-* cells were included in the subsequent analysis (Fig 3B). Both aging and IL7RKO saw an unbalanced expansion of two distinct cell populations (Fig 3B, Supplemental fig 6A-E). In the case of IL7RKO, the NC subset was drastically expanded in some samples (Fig 3B, Supplemental fig 6A-C,). As this cluster expresses high levels of cKit (Supplemental fig 6C) the observation was readily confirmed by flow cytometry (Supplemental fig 6D). In aging on the other hand, a new population of cells (oSMC) expressing smooth muscle cell markers along with osteogenic marker *Sost* appeared in some samples, suggesting a possible connection to vessel calcification seen in aged individuals ^40^ (Fig 3B, Supplemental fig 6A-B, E). We also observed a notable expansion of endothelium and pericytes in aging samples (Fig 3B, Supplemental fig 6A-B).

**Figure 3.**
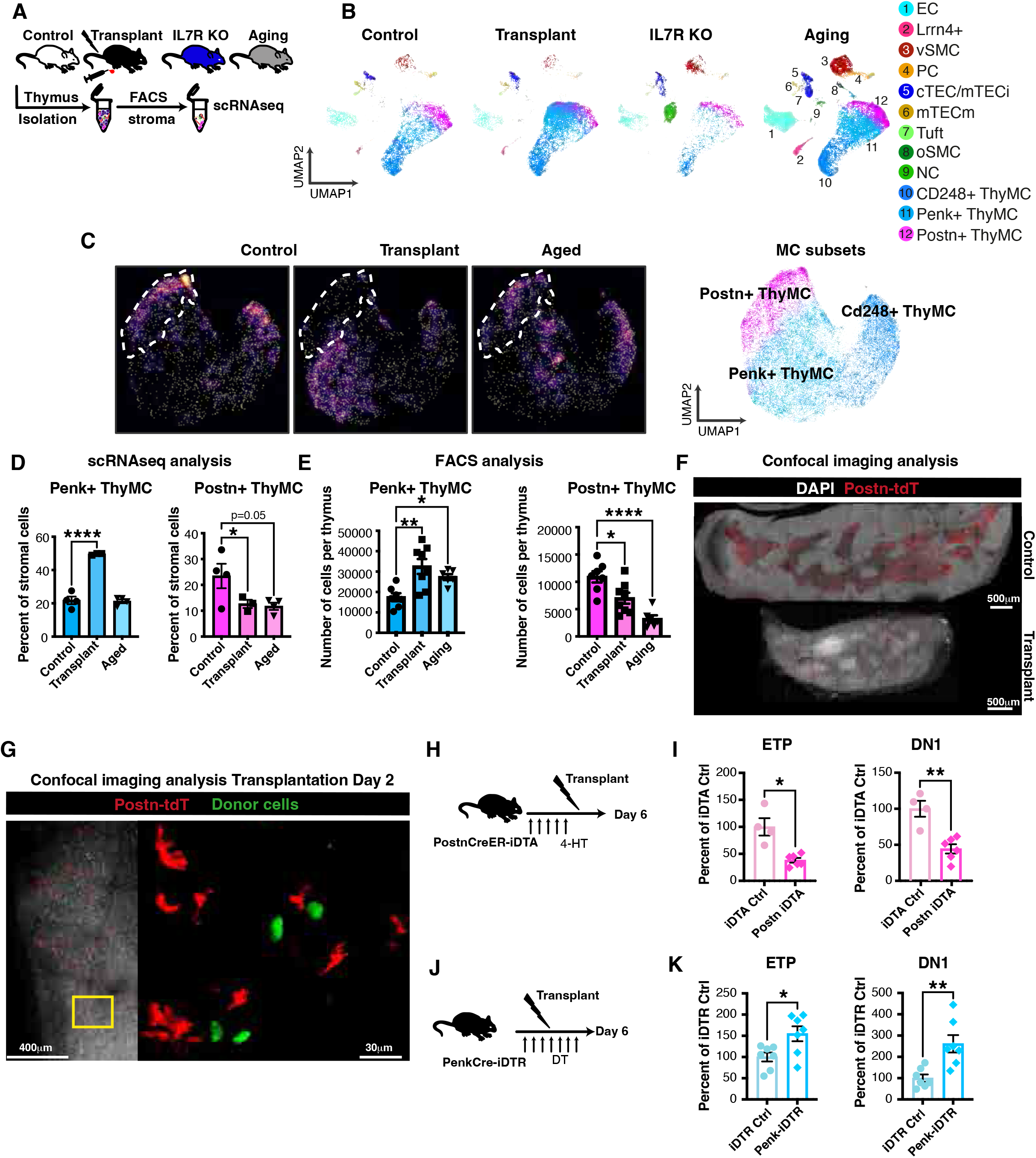
Postn+ ThyMCs are lost during transplant conditioning and aging (A) Schematic overview of sample collection from Transplantation, IL7RKO and Aging stress conditions. (B) UMAP embeddings showing detailed annotation of different stromal cell populations (*Cd3e-, Cd4-, Cd8a-, Ptprc-)* in Control (n=4, total number of stromal cells=5451), Transplant (n=3, total number of stromal cells=8057), IL7RKO (n=4, total number of stromal cells=4551) and Aging (n=4, total number of stromal cells=16178) samples. (C) The difference between Control and Transplantation, and Aging samples visualized via condition-specific densities in a joint ThyMC specific UMAP embedding (left). Detailed annotation of ThyMC subsets shown in ThyMC specific UMAP embedding (right). Control (n=4), Transplantation (n=3), and Aging (n=4), total number of ThyMC = 19739. (D) Bar graphs comparing proportional shifts of individual ThyMC subsets between Control, Transplant, and Aging samples as determined by scRNAseq. (Control n= 4, Transplant n= 3, Aging n=4) Statistical significance is based on beta regression with Benjamini- Hochberg FDR control of these comparisons. (E) Bar graphs showing flow cytometric quantification of absolute numbers of different ThyMC subsets in Control, Transplant (day 3) or Aged (2 years old) *Penk-Cre-tdTomato* mice. Statistical significance was calculated using one-way ANOVA followed by Dunnet’s post-hoc analysis. (F) 3D confocal microscopy images from Control and Day 3 post-transplant *Postn-CreER- tdTomato* thymi stained with DAPI (white; cell nuclei), and tdTomato (red; Postn+ ThyMCs). (G) 3D confocal microscopy images of thymic tissue from an irradiated *Postn-CreER- tdTomato* mouse transplanted with 40 000 GFP+ lymphoid progenitors cells 2 days post- transplantation. Tissue was stained with DAPI (white; cell nuclei), tdTomato (red; Postn+ ThyMCs) and GFP (green; lymphoid progenitor donor cells). (H) Schematic illustration of experimental design for depletion of Postn+ cells in *Postn- CreER-tdT/iDTA* mice. (I) Flow cytometric quantification of early thymic progenitors and DN1 thymocytes in Control (iDTA Ctrl n=4) and *Postn-CreER-tdT/iDTA* (Postn iDTA n=6) mice 6 days post-bone marrow transplantation. Statistical significance was calculated using unpaired two-tailed student’s t-test. Two independent experiments. (J) Schematic illustration of experimental design for depletion of Penk+ cells in *Penk-Cre- tdT/iDTR* mice. (K) FACS analysis of early thymic progenitors and DN1 thymocytes in Control (iDTR Ctrl n=7) and *Penk-Cre-tdT/iDTA* (Penk iDTA n=7) mice 6 days post-bone marrow transplantation. Statistical significance was calculated using unpaired two-tailed student’s t-test. Two independent experiments.

The ThyMC compartment undergoes significant remodeling across all three types of stress states. In HSCT samples Penk+ ThyMCs were found to be strikingly increased and the T cell supportive Postn+ ThyMC were significantly reduced (Fig 3B-D). Aging samples display a similar trend of Postn+ ThyMCs loss (Fig 3B-D) and within the ThyMC population there was an evident bias toward Penk+ ThyMCs (Supplemental fig 6F). These observations were further validated flow cytometric analysis of *Penk-tdT* HSCT recipients and 2-year old reporter mice (Fig 3E). Tissue-wide confocal imaging of *Postn-CreER-tdT* thymi 3 days post-bone marrow transplantation further confirmed Postn+ ThyMC depletion (Fig 3F, Supplemental fig 6G). Lastly, sorted CD248- ThyMCs from aged mice expressed significantly lower levels of *Postn* and *Ccl19* compared to young controls further confirming that Postn+ ThyMCs are diminished in aging (Supplemental fig 6H). Interestingly, the total ThyMC population was also found to be significantly depleted in IL7RKO as well as aged animals (Supplemental fig 6A, I). As both IL7RKO and aging are conditions characterized by prolonged reduction in T cell development, this is in line with previous observations that maintenance of ThyMCs depend on interactions with maturing lymphocytes ^26^. Taken together, these data demonstrate that Penk+ ThyMCs respond to both the acute injury of transplantation and the chronic stress of aging by expanding, whereas the T cell supportive Postn+ ThyMCs are depleted under the same conditions.

GSEA on genes differentially expressed between stress conditions demonstrated that similar pathways were upregulated across all three ThyMCs subsets. In transplantation samples all ThyMCs were enriched in inflammatory stress terms and adipogenic pathways (Supplemental fig 6J). Aging similarly induced pro-adipogenic pathways across ThyMCs but also an increase in ECM deposition (Supplemental fig 6J) consistent with the increased fat and fibrosis seen in the aging thymus ^41,42^. Importantly, T cell terms were significantly down regulated in all ThyMC subsets and this was further supported by assessment of Ccl19 and Flt3l protein levels in whole thymic tissue 3 days post-transplant (Supplemental fig 6K).

Levels of both cytokines were significantly lower in transplant tissue, indicating that other stromal populations were not able to make up for the Postn+ ThyMC depletion (Supplemental fig 6K).

Because Postn+ ThyMCs express factors that would act on early steps in T cell development ^35-37^ we assessed if Postn+ cells and thymic seeding progenitors are localized in the same region, 4-HT induced *Postn-CreER-tdT* were radiation conditioned and transplanted with 40 000 GFP labeled lymphoid progenitor cells (LPC, lineage-cKit+CD135+). The thymic tissue was imaged two days later, demonstrating relative proximity of newly arrived GFP+ LPCs with areas rich in tdT+ Postn+ ThyMCs (Fig 3G).

To examine the functionality of the ThyMC subsets we intercrossed *Penk-Cre* and *Postn- CreER* mice with deleter lines *Rosa26-LSL-iDTR* (*iDTR*) or *Rosa26-LSL-DTA* (*iDTA*) respectively. Selective, efficient depletion (Supplemental fig 7A) of Postn+ cells was induced just prior to radiation conditioning and bone marrow transplant (Fig 3H). Isolation of thymi 6 days later revealed an expected decrease in CD248- ThyMCs and a concomitant increase of CD248+ ThyMCs but no alterations of other stromal populations (Supplemental fig 7B)

While later maturation stages remained unchanged at this time point, deletion of Postn+ cells resulted in a drastic reduction of the earliest steps in T cell development; ETP and DN1 (Fig 3I, Supplemental fig 7C). To confirm that the effects of Postn+ cell depletion were thymus specific, we induced cell deletion *in vitro* by culturing adult thymic organ cultures (ATOCs) from *Postn-CreER-tdT/iDTA* mice for 8 days in the presence of 4-HT at which point no Postn+ cells could be detected (Supplemental fig 7D). Tissue-wide 3D confocal imaging confirmed that stromal structures relevant to T cell development remained after 8 days of culture (Supplemental fig 7E). *Postn-CreER-tdT/iDTA* ATOCs contained fewer overall ThyMCs but all other stromal cell types were unaltered (Supplemental fig 7G). The absolute number ETPs and DN1 thymocytes were significantly reduced compared to controls (Supplemental fig 7F). This demonstrates that loss of Postn+ ThyMCs directly effects early steps of T cell development.

For assessment of the long-term effects of Postn+ ThyMC depletion, control or deleted *Postn-CreER-tdT/iDTA* ATOCs were grafted under the kidney capsule of athymic *Foxn1*^*null/null*^ mice (Supplemental fig 8A). Blood sampling 8 weeks post-implantation showed significantly fewer T cells in circulation in *Foxn1*^*null/null*^ mice that received a Postn+ ThyMC depleted graft compared to *iDTA* control samples (Supplemental fig 8B). In addition, sjTREC analysis of the implanted ATOCs demonstrated that *de novo* synthesis of T cells was impaired in response to Postn+ ThyMC depletion 10 weeks post-grafting (Supplemental fig 8C). This suggests that Postn+ ThyMCs are essential for the first steps of T cell development and ultimately for long-term upkeep of T cell output.

Ablation of Penk+ cells (Supplemental fig 8D) was tested in the context of radiation conditioning and HSCT by injection of diphtheria toxin in *Penk-Cre-tdT/iDTR* mice (Fig 3J). Loss of Penk+ cells in bone marrow transplant recipients 6 days post-procedure did not alter the thymic endothelium or epithelium, it did however result in a significant reduction of both CD248- and CD248+ ThyMC (Supplemental fig 8E). Interestingly, selective deletion of Penk+ cells had the opposite effect to the absence Postn+ cells (Fig 3I), the first steps of T cell development showed significant improvements (Fig 3K) and no effects on more mature stages (Supplemental fig 8F). *In* vitro deletion in *Penk-Cre-tdT/iDTR* ATOCs confirmed this by a significant reduction in CD248- ThyMCs but no changes in other stromal subsets (Supplemental fig 8G) and increased numbers of ETPs as well as DN1 thymocytes (Supplemental fig 8H). This indicates that the increase of Penk+ ThyMCs seen under the stress of transplantation and aging confers an inhibitory effect on early T cell development.

### Postn+ ThyMCs recruit T cell progenitors and regenerate T cell immunity

Since Postn+ ThyMC are lost upon transplant conditioning we tested whether replenishing T cell supportive ThyMCs specifically could enhance thymic regeneration. *Penk-Cre* mice were crossed with *Rosa26-mT-LSL-mG* (*mTmG*) mice to enable fluorescent tracing of both Cre- cells (Postn+ cells; tdTomato+) and Cre+ cells (Penk+; GFP+). Subsequent isolation and intrathymic transfer of Penk+ or Postn+ ThyMCs in the context of HSCT (Fig 4A), demonstrated that Postn+ (tdTomato+) as well as Penk+ (GFP+) ThyMCs persisted in the tissue six days later (Supplemental fig 9A). Recipients of Postn+ ThyMCs had improved ETP and endothelial cell numbers after (Fig 4B, Supplemental fig 9B, Supplementary table 2) compared to PBS Sham, or Penk+ ThyMC treated mice. These data align with the enrichment in GO terms of T cell development and angiogenesis (Fig 2C). Other stromal cell types were unaltered (Supplemental fig 9B, Supplementary table 2). Transfer of Postn+ ThyMCs thus improved short-term thymic regeneration through recruitment of T competent progenitors and preservation of endothelium.

**Figure 4.**
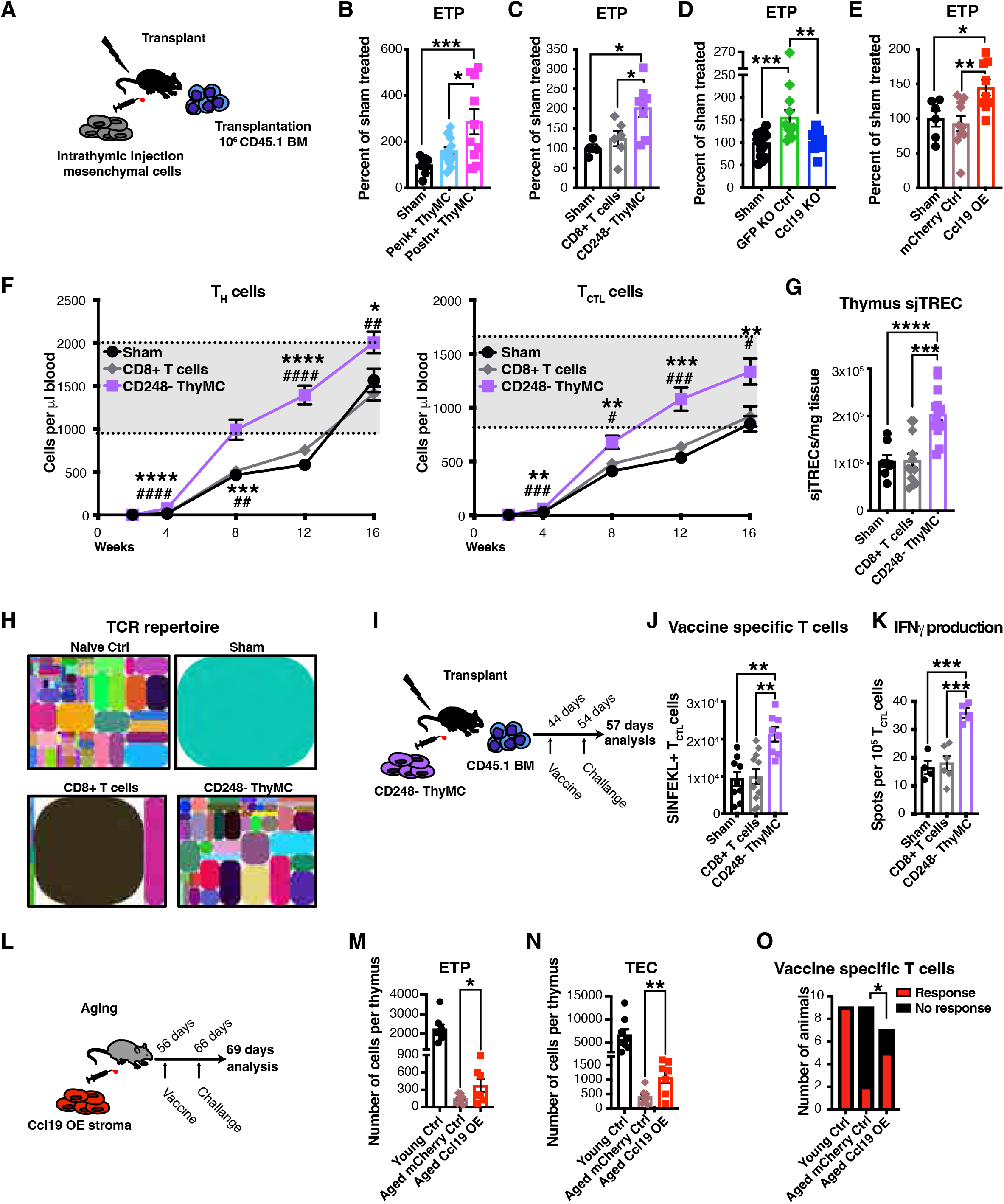
Postn+ ThyMCs maintain and recruit T cell <u>progenitors</u> and regenerate T cell immunity (E) Schematic illustration of experimental design for adoptive transfer of mesenchymal cells in the context of HSCT. (E) FACS quantification of early thymic progenitors after Sham (PBS), GFP+ (Penk+) ThyMC or tdTomato+ (Postn+) ThyMC treatment displayed as bar graphs. (Dose of Penk+ and Postn+ ThyMCs: 4000-8000) (Sham=9, Penk+ ThyMC=14, Postn+ ThyMCs=10) Three independent experiments. Statistical significance was determined by one-way ANOVA followed by Tukey’s post-hoc analysis. (E) Bar graphs displaying the flow cytometric quantification of early thymic progenitors 6 days following an intrathymic sham injection or adoptive transfer of CD8+ T cells and CD248- ThyMCs in HSCT recipients. (Dose of CD8+ T cells and CD248- ThyMCs: 2000-4000) Values are presented as percent of Sham treated animals and statistical significance was assessed using one-way ANOVA followed by Tukey’s post-hoc analysis (Sham n= 4, CD8+ T cells n=6, CD248- ThyMC n=8). Two independent experiments. (E) Bar graphs showing the results of FACS quantification of early thymic progenitors 6 days after an intrathymic sham injection or adoptive transfer of Cas9-GFP CD248- ThyMCs following knockout of either GFP control (KO Ctrl) or Ccl19 (KO) in mice undergoing HSCT. (Dose of GFP KO Ctrl and Ccl19 KO cells: 4000) Values are presented as percent of Sham treated animals and statistical significance was assessed using one-way ANOVA followed by Tukey’s post-hoc analysis (n= 15 Sham, n=11 GFP KO Ctrl, n=10 Ccl19 KO). Three independent experiments. (E) Bar graphs showing the results of FACS analysis of early thymic progenitors and endothelial cells, 6 days following an intrathymic sham injection or adoptive transfer of Cas9-GFP bone marrow stromal cells following infection with of either mCherry control (Ctrl) or Ccl19 mCherry overexpression (Ccl19 OE) vectors in HSCT recipients. (Dose of mCherry Ctrl and Ccl19 OE cells: 50 000) Values are presented as percent of Sham treated animals and statistical significance was assessed using one-way ANOVA followed by Tukey’s post-hoc analysis (n= 6 Sham, n=9 mCherry Ctrl, n=9 Ccl19 OE). Two independent experiments. (E) Flow cytometric analysis of cytotoxic T lymphocytes (T_CTL_ cells) and T helper cells (T_H_ cells) in peripheral blood following adoptive transfer of Sham, CD8+ T cells or CD248- ThyMCs 2-16 weeks post-bone marrow transplantation. (Dose of CD8+ T cells and CD248- ThyMCs: 10 000) (Sham=7, CD8+ T cells=9, CD248- ThyMCs=9) Two independent experiments. Statistical significance was determined by one-way ANOVA followed by Tukey’s post-hoc analysis. # denotes statistical significance for CD248- ThyMC vs CD8+ T cell comparison and * denotes statistical significance for CD248- ThyMC vs Sham comparison. Gray shaded area denotes the peripheral blood parameters of untreated control mice (n=12) housed in the animal facility at the same time as the transplant recipients. (E) Quantification of *de novo* generated T cells following adoptive transfer of Sham, CD8+ T cells or CD248- ThyMCs by way of signal-joint T cell receptor excision circles (sjTRECs) 4 weeks following bone marrow transplantation. (Dose of CD8+ T cells and CD248- ThyMCs: 10 000) (Sham=8, CD8+ T cells=12, CD248- ThyMCs= 14) Two independent experiments. Statistical significance was determined by one-way ANOVA followed by Tukey’s post-hoc analysis. (E) Diversity in T cell receptors (TCRs) as analyzed using the sequencing of the CDR3 β chain 4 weeks following adoptive transfer of Sham, CD8+ T cells or CD248- ThyMCs and HSCT. (Dose of CD8+ T cells and CD248- ThyMCs: 10 000) Each colored square represents a single clone. Samples were pooled from sorted CD3+ splenocytes from four mice for each group and the combined data is represented. (E) Overview of ovalbumin vaccination response assessment after adoptive transfer of CD248- ThyMCs in HSCT recipients. (E) Bar graphs showing the flow cytometric quantification of absolute numbers of ovalbumin specific T_CTL_ cells in recipients of CD248- ThyMCs and HSCT. (Dose of CD8+ T cells and CD248- ThyMCs: 10 000) (Sham=4-9, CD8+ T cells=6-11, CD248- ThyMCs= 4-8) Statistical significance was determined by one-way ANOVA followed by Tukey’s post- hoc analysis. Two independent experiments. (E) Bar graph representation of IFNg production by ovalbumin specific T_CTL_ cells as assessed by ELISpot. (Dose of CD8+ T cells and CD248- ThyMCs: 10 000) (Sham=4-9, CD8+ T cells=6-11, CD248- ThyMCs= 4-8) Statistical significance was determined by one-way ANOVA followed by Tukey’s post-hoc analysis. (E) Schematic depiction of ovalbumin vaccination experimental design for aged recipients of *Ccl19* overexpressing bone marrow stroma. (E) Bar graphs showing the results of flow cytometric analysis of ETPs in Young controls (Ctrl), Aged mCherry controls (Ctrl) and Aged Ccl19 mCherry overexpression (Ccl19 OE) recipients. (Dose of mCherry Ctrl and Ccl19 OE cells: 150 000) Statistical significance was determined using Student’s t-test comparing Aged mCherry Ctrl with Aged Ccl19 OE. (Young n= 9, Aged mCherry Ctrl n=9, Aged Ccl19 OE n=7). Two independent experiments. (E) Bar graphs representing the flow cytometric analysis of TECs in Young Ctrl, Aged mCherry Ctrl and Aged Ccl19 OE treated mice. (Dose of mCherry Ctrl and Ccl19 OE cells: 150 000) Statistical significance was determined using Student’s t-test comparing Aged mCherry Ctrl with Aged Ccl19 OE. (Young n= 9, Aged mCherry Ctrl n=9, Aged Ccl19 OE n=7). Two independent experiments. (E) Bar graphs showing the number of animals that failed or succeeded in mounting an ovalbumin specific CD8+ T_CTL_ cell response as assessed by flow cytometric analysis of SINFEKL labeling, after an intrathymic injection of bone marrow stromal cells overexpressing either mCherry Ctrl or Ccl19 OE. (Dose of mCherry Ctrl and Ccl19 OE cells: 150 000) Statistical significance was determined using Student’s t-test comparing Aged mCherry Ctrl with Aged Ccl19 OE. (Young n= 9, Aged mCherry Ctrl n=9, Aged mCherry Ccl19 OE n=7). Two independent experiments.

Next, CD248- ThyMCs (Supplemental fig 4A and G) from *Ubiquitin-GFP* mice were injected intrathymically into irradiated HSCT recipients (Fig 4A). As control, syngeneic single-positive CD8 thymocytes, which are clearly distinct from mesenchymal cells, were intrathymically injected. Analysis six days post-transplantation, demonstrated GFP-labeled CD248- ThyMCs persist in the thymus (Supplemental fig 9C). CD248- ThyMC treated mice had increased ETPs and endothelial cells in comparison to Sham and CD8+ T cell treated animals (Fig 4C, Supplemental fig 9D, Supplementary table 2). ThyMCs and TEC numbers (Supplemental fig 9D) remained unchanged. As the CD248- population is a mixture of Penk+ ThyMCs and Postn+ ThyMCs these data confirmed that even this heterogeneous ThyMC graft is capable of enhancing thymus regeneration.

One of the factors preferentially expressed by thymic ThyMCs is *Ccl19* (Fig 2D). As a known ligand of Ccr7 that is necessary for thymic homing, Ccl19 has been indirectly implicated in thymic development and ETP recruitment ^36^. To determine if *Ccl19* expression in ThyMCs was necessary for the observed improvement in ETP seeding after transplantation, CD248- ThyMCs were isolated from Cas9-GFP expressing mice. The cells were transduced with lentiviral vectors expressing *Ccl19* or control *GFP* gRNAs (knockout efficiency ∼70%) and the *Ccl19* knockout cells were confirmed to have a reduced ability to attract CCR7+ thymocytes *in vitro* (Supplemental fig 9E). Intrathymic transplantation (Fig 4A) demonstrated that *Ccl19* knockout abrogated the improvement in ETP recruitment following CD248- ThyMC injection (Fig 4D, Supplementary table 2) compared to *GFP* targeted controls. Regeneration of stromal cells was not impacted by *Ccl19* deletion (Supplemental fig 9F). We further tested the importance of *Ccl19* expression by ectopically expressing it in primary bone marrow stromal cells. Low passage, primary bone marrow stromal cells were infected with a *mCherry* control or *mCherry-Ccl19* overexpression vector and confirmed to express functional Ccl19 protein (Supplemental fig 10H). The bone marrow stroma was transferred intrathymically to HSCT recipients (Fig 4A) and flow cytometry 6 days later demonstrated persistence of rare mCherry+ cells (Supplemental fig 9G). Mice that received an intrathymic injection of *mCherry Ccl19* cells had increased ETPs compared to Sham treated or *mCherry* controls (Fig 4E, Supplementary table 2) but no changes in their stromal cell subsets (Supplemental fig 9I). Consequently, *Ccl19* is both necessary and sufficient for the ability of ThyMC’s to recruit ETPs to the thymus.

To determine if the increased influx of progenitors resulted in long-term improvements in T cell number following HSCT, recipients of Sham, CD8+ T cells or CD248- ThyMC treatments were followed for 16-weeks post-transplant. Analysis of peripheral blood demonstrated dramatically increased numbers of CD4+ T_H_ cells and CD8+ T_CTL_ cells in CD248- ThyMC recipients (Fig 4F), with no impact on B cells or myeloid populations (Supplemental fig 10A). Notably, GFP+ ThyMCs were still detectable 16 weeks after transplantation (Supplemental fig 10B). Therefore CD248- ThyMCs are able to engraft and provide durable enhancement of T cell regeneration following acute thymic injury and HSCT.

In order to ensure that the improved T cell reconstitution was due to *de novo* generation of T cells, sjTREC analysis was performed on thymi 4 weeks following intrathymic injection of CD248- ThyMCs (Fig 4A). Thymus cellularity was significantly higher in ThyMC treated mice (Supplemental fig 10C). The sjTREC analysis demonstrated that production of newly rearranged T cells was significantly improved in the mice injected with CD248- ThyMCs compared to Sham and CD8+ T cell treated controls (Fig 4G). In line with this T cell receptor (TCR) sequencing revealed greater diversity in CD248- ThyMC treated HSCT recipients (Fig 4H).

To test whether functional immunity was affected, transplantation recipients were vaccinated against the neoantigen ovalbumin after 44 days (Fig 4I). CD248- ThyMC treated mice had significantly improved immune responses as evidenced by increased numbers of ovalbumin specific CD8+ T_CTL_ cells producing IFNg in response to ovalbumin exposure (Fig 4J-K, Supplemental fig 10D). Thus, the improvements in early thymic regeneration seen after CD248- ThyMC transfer ultimately translated into a robust production of T cells and T cell immunity. Thymic ThyMCs represent a novel T cell regenerative stromal cell type in the thymus atrophic after acute injury.

To test whether a similar approach can be useful in the context of chronic age-related thymic atrophy, we used a model that has potential for clinical translation: enforced *Ccl19* expression in cultured bone marrow derived ThyMCs. 24-month old male and female mice were intrathymically injected with *mCherry* control or *mCherry-Ccl19* overexpressing bone marrow stromal cells and subsequently vaccinated against ovalbumin 8 weeks later (Fig 4L). Flow cytometric analysis of the thymic tissue from these mice confirmed persistence of *mCherry*+ cells within the stromal compartment at 9.5 weeks (Supplemental fig 10F). This was accompanied by a significant increase in the number of ETPs, TECs and total thymocytes in the aged mice that received *mCherry-Ccl19* overexpressing stroma compared to the *mCherry* controls (Fig 4M-N, Supplemental fig 10E). Vaccinated, young control mice responded promptly by expanding ovalbumin specific (SINFEKL+) CD8+ T_CTL_ clones (Fig 4O). As expected, the vaccine response in the aged *mCherry* controls was severely impaired, with only 2 of 9 mice mounting detectable levels of ovalbumin specific CD8+ T_CTL_ cells (Fig 4O). Notably, in *mCherry-Ccl19* recipients, the majority of the mice (5 of 7) responded to the vaccine (p= 0.0024) (Fig 4O) and the mice that responded generated equivalent numbers of (SINFEKL+) CD8+ T_CTL_ cells as young mice (Supplemental fig 10G). Taken together these results demonstrate that Ccl19+ stromal cells are sufficient to induce rejuvenation in the aged thymus and regenerate T cell vaccine responses.

## Discussion

In every tissue type, the stromal compartment has a role in governing normal tissue development with a growing appreciation of functions in adult tissue homeostasis and regeneration. Heterogeneity among the stromal populations is increasingly well annotated in hematopoietic tissue ^27,28,43^ with delineation of specific cell types in parenchymal stem cell support, tissue regeneration and even tissue malignant transformation now experimentally defined ^44-46^. Targeting stromal cells pharmacologically to alter tissue outcomes has benefit in some settings of regeneration after injury ^15,47,48^. Adoptive transfer of stromal cells has been less explored though infusing endothelial cells has been shown to enhance over all hematopoietic recovery as well as T cell regeneration specifically ^15,49^. Here we define three functionally distinct subsets of ThyMCs in both mouse and human thymus. The lineage relationship between them is unclear, but they have demonstrably different roles particularly in recruiting hematopoietic cells via Ccl19. Even modest engraftment of Ccl19+- mesenchymal cells increased early thymic progenitors and led to improved mature T cell generation. This is consistent with earlier work demonstrating that an increase in early T cell precursors improves overall thymic regeneration ^50^ suggesting that homeostatic feedback within the thymus amplifies the effects of stromal manipulation to achieve substantial effects on immune regeneration. Notably, while mesenchymal populations have been used as treatment in many settings, the durable engraftment of the cells that we observe at 16 weeks and the specific subset of cells that provide benefit suggest that this is a novel mesenchymal population and application. Our data indicate that specific subsets of thymic mesenchymal cells may improve a critical and currently unmet medical need of thymic regeneration.

## Material and Methods

### Animals

Sequencing and transplantation experiments involving young mice were carried out using 8 weeks old male and female C57Bl/6J mice or B6.129S7-Il7rtm1Imx/J. For experiments involving aged animals 90 weeks old male and female C57Bl/6J mice were purchased from Jackson laboratories and allowed to age an additional 3 months before being used for experiments. B6.SJL-Ptprca Pepcb/BoyJ (CD45.1) and C57BL/6-Tg(UBC-GFP)30Scha/J mice were used as donors for HSCTs. B6.Cg-Foxn1nu/J animals served as recipients for kidney capsule grafting. B6.Cg-Gt(ROSA)26Sortm9(CAG-tdTomato)Hze/J (*tdT*) mice were crossed with B6;129S-Penktm2(cre)Hze/J (*Penk-Cre)* and B6.129S- Postntm2.1(cre/Esr1*)Jmol/J (*Postn-CreER*) mice respectively to generate mesenchymal cell subset specific reporters. Genetic deletion of Postn+ ThyMC was achieved by breeding B6.129S-Postntm2.1(cre/Esr1*)Jmol/J with B6.129P2-Gt(ROSA)26Sortm1(DTA)Lky/J (*iDTA*) animals. Specific depletion of Penk+ ThyMC was instead done by crossing B6;129S- Penktm2(cre)Hze/J mice with the C57BL/6-Gt(ROSA)26Sortm1(HBEGF)Awai/J (*iDTR*) deleter line. Donors for adoptive transfer of Penk+ and Postn+ ThyMC were generated by crossing B6.129(Cg)-Gt(ROSA)26Sortm4(ACTB-tdTomato,-EGFP)Luo/J (*mTmG*) reporter mice with the B6;129S-Penktm2(cre)Hze/J Cre line. To enable deletion of individual factors in thymic ThyMCs, cells were isolated from B6J.129(B6N)-Gt(ROSA)26Sortm1(CAG- cas9*,-EGFP)Fezh/J animals. All mice were obtained from Jackson Laboratories and all animal experimentation was carried out in accordance with national and institutional guidelines and with the approval of the Institutional Animal Care and Use Committee of Massachusetts General Hospital.

### Tissue collection and processing

All human tissue specimens were collected with institutional review board (IRB) approval. The tissue was processed immediately upon isolation to ensure highest possible cell quality. Murine samples were cut into fine pieces and digested in Medium 199 (M199, Gibco) with 2% (v/v) fetal bovine serum (FBS, Gibco), Liberase (0.5WU/ml, Roche) and DNAse I (0.1 KU, ThermoFisher) 3 × 15 minutes at 37°C under constant agitation. Human samples were processed by digestion with M199 with 2% FBS, DNAse I (0.1 KU) and 2 mg/ml Stemxyme 1 (Worthington) for 2 × 30 minutes at 37°C under constant agitation. For the last 30 minutes the samples were digested with the Stemxyme/DNAse I cocktail in combination with 0.125% Trypsin (Gibco). All samples were digested in the presence of RNase inhibitors (RNasin (Promega) and RNase OUT (Invitrogen).

Primary bone marrow stromal cells were collected from tibias and femurs by crushing in 2% (v/v) FBS in PBS and the resulting bone fragments were repeatedly washed to remove hematopoietic contaminants. The bone was thereafter finely cut and digested with 0.25% Collagenase I (Stem Cell Technologies) and DNAse I (25 Units/ml) for 60 minutes at 37°C under constant agitation. The cells released cells were subsequently cultured under hypoxic conditions (2% O_2_) in α-MEM (ThermoFisher) supplemented with 20% FBS (Gibco), and 1% Penicillin/Streptomycin (Gibco).

### FACS sorting for single-cell RNA sequencing

After blocking with anti-human CD16/32 Fc-block (BD Biosciences) for 10 minutes at 4°C, human single cell suspensions were stained with Lineage cocktail-FITC, CD66b-FITC, CD45-BV711, CD235a-BV711, CD8a-APC/Cy7 and CD4-BV605 (all from BD Biosciences). Mouse samples were also blocked with anti-mouse CD16/32 Fc-block (BD Biosciences) for 10 minutes at 4°C, followed by staining with CD45-PE/Cy7 and Ter119-PE (Both from BioLegend). Samples were stained for 45 minutes at 4°C under constant agitation. For detection of dead cells 7-AAD (ThermoFisher) was added to the samples immediately before analysis. Flow sorting for live and non-hematopoietic cells (7-AAD, CD45-CD235a/Ter119- Lineage-) was performed on a BD FACS Aria III equipped with a 70um nozzle (BD Biosciences).

### Single-cell RNA sequencing

Sorted thymus stromal cells were encapsulated into emulsion droplets using the Chromium Controller (10X Genomics). scRNA sequencing libraries were subsequently prepared using Chromium Single Cell 3 v2 and v3 Reagent kits (10X Genomics). Libraries were diluted to 4nM and pooled before sequencing on the NextSeq 500 Sequencing system (Illumina).

### scRNA seq analysis

Single-cell RNA sequencing data was processed using the CellRanger suite (v. 6.1.2). For human samples, the GRCh38 reference (GENCODE v32/Ensembl 98) was used. Mouse samples were aligned to mm10 (GENCODE vM23/Ensembl 98) reference. Standard CellRanger filtration strategy was used for empty droplet identification. Additionally, cells with more than 10% mitochondrial UMI counts and 15,000 total UMIs (25,000 for samples that were obtained using 10x v3 chemistry) were also removed (fig. S1C). Doublet detection was performed with Scrublet (v. 0.2.2) ^51^.

Data was normalized using the pagoda2 pipeline. Firstly, data was log(x + 1)- transformed. Secondly, top-3000 overdispersed genes in each sample were selected for PCA. Finally, all samples were integrated together using conos (v. 1.4.3) ^52^ function *buildGraph* based on the first 30 PCs from each sample. Downstream analysis and data visualization was performed using the SCANPY (v. 1.7.2) ^53^ Python package. The Leiden (v. 0.8.4) ^54^ algorithm was used for unsupervised clustering of the data. High-level cell type identification was performed on common embedding by examining expression of literature derived markers. Mesenchymal cells subtype identification was performed on the novel embedding that was created by re-normalization, re-integration and re-clustering of only ThyMCs (in this case top- 1000 overdispersed genes and first 15 PCs were used in the integration pipeline), following the procedures described above. The ThyMCs subpopulations were defined by unbiased Leiden clusters, and named based on the detected marker genes. Embedding density was computed and visualized with Cacoa [https://doi.org/10.1101/2022.03.15.484475] R package (v. 0.1).

Markers of cell populations were identified with Mann-Whitney U-test and Benjamini-Hochberg FDR control on log-normalized data. Differential expression analysis for evaluation of inter-sample expression variability was performed with the DESeq2 R package ^55^ on pseudo- bulk counts. GSEA analysis with GSEApy ^56,57^ was performed on the ranked list of differentially expressed genes. Only Gene Ontology ^58^ gene sets were used in this analysis. Human and mouse identifiers were mapped using bioMart to human identifiers for merging of datasets between species. Data was re-processed and analyzed jointly as described below.

### 4-hydroxytamoxifen injections

In order to induce expression of Cre in B6.129S-Postntm2.1(cre/Esr1*)Jmol/J mice 4- hydroxytamoxifen (4-HT) (Selleck Chemical) was dissolved at 40 mg/ml in absolute ethanol by heating at 56°C with 250rpm shaking. The 40 mg/ml 4-HT stock solution was diluted 1:1 in Kolliphor EL (MilliporeSigma) ^59^.Before injection the solution was further diluted 1:3 in PBS (Gibco) and warmed to 37°C. Mice were injected intraperitoneally with 200 μg/g mouse for 5 consecutive days. Depletion efficiency was determined by flow cytometry 2 days after the final injection.

### Diphtheria toxin injections

Selective depletion of Penk+ cells was achieved by injecting *Penk-Cre-tdT/iDTR* mice with diphtheria toxin (MilliporeSigma). In brief mice were intraperitoneally injected with 25 ng/ diphtheria toxin dissolved in PBS for 7 consecutive days ^60^. Depletion efficiency was determined by flow cytometry 2 days after the final injection.

### Intracellular staining for cytokines and FACS analysis of human thymus stroma

To analyze the major thymic stromal cell types in human samples single cells suspensions non-specific antibody binding was blocked with human Fc-Block (BD Bioscience) and the cells were subsequently stained with CD45-PE, CD235a-PE, CD4-BV605 CD31- PE/Dazzle594, CD326-BV421 (all from Biolegend) and CD8-APC/Cy7 (BD Biosciences). The samples were stained with LIVE/DEAD Fixable Blue dead cell stain (Thermo Fischer Scientific) to enable exclusion of dead cells. Subsequently, the samples were fixed and permeabilized using BD Bioscience Fixation/Permeabilization Solution Kit in accordance with the instructions supplied by the manufacturer. Lastly the samples were incubated with IL-15-AlexaFluor700 (R&D Systems), Ccl19-AlexaFluor488 (Bioss) and Flt3lg-AF647 (Abcam). The latter antibody was conjugated in house the Apex AlexaFluor647 antibody labeling kit from Thermo Fischer Scientific.

### FACS sorting and analysis of thymus stromal cell populations

For analysis of various thymus stromal cell populations human samples were blocked with human Fc-block (BD Bioscience) for 10 minutes at 4°C and then stained with Lineage cocktail-FITC, CD66b-FITC, CD45-BV711, CD235a-BV711, CD8a-APC/Cy7 and CD4- BV605 in combination with CD326-BV421 (BD Bioscience) and CD31-PE/Dazzle594 (BioLegend). Murine stromal cell types were characterized and sorted by surface staining for CD45-APC/Cy7, Ter119-APC/Cy7 and CD11b-BV421 (BD Biosciences), Dpp4-BV421, Pdpn-AF488, F4/80-AF700 (Biolegend), Ltbr-Biotin (ThermoFisher) as well as CD31- BUV737, CD326-BV711, and CD140a-BV785 (all from BD Biosciences). Itgb5, CD99l2 and CD248 (R&D Systems) were conjugated in house to PE/Cy7 and APC (Abcam) respectively and also used for some of the stromal cell sorts. For panels requiring streptavidin labeling cells were stained with streptavidin-BV570 (Biolegend). Intracellular staining for Cre (Cre-AF488 Novus Biologicals) and S100A4 (S100A4-AF700 Novus Biologicals), was carried out using BD Bioscience Fixation/Permeabilization Solution Kit in accordance with the instructions supplied by the manufacturer. Flow cytometric analysis was performed on a BD FACS Aria III (BD Biosciences).

### FACS analysis of peripheral blood chimerism

Transplantation recipients were bled retro-orbitally under isoflurane anesthesia and using heparinized capillaries and blood was collected into EDTA containing tubes to prevent coagulation. Complete blood counts were obtained using the Element Ht5 Auto Hematology analyzer. RBCs were subsequently lysed, and cells were blocked with murine Fc-block (BD Biosciences) in in PBS with 2% (v/v) FBS for 10 minutes at 4°C. The samples were then stained using the following antibodies: CD11b-PE/Cy7, Gr-1-PE/Cy7, (ThermoFisher), CD45.1-AF700, CD4-APC, CD3e-PE, CD8a-BV570, CD19-BV650 (BioLegend), CD45.2- BUV395 and B220-BV650 (BD Biosciences). Flow cytometric analysis was performed on a BD FACS Aria III (BD Biosciences).

### FACS analysis of thymocyte subsets

Thymic hematopoietic cells were obtained by gentle mechanic dissociation of the tissue. For quantification of the different developmental steps in T cell maturation the cells were blocked with murine Fc-block (BD Biosciences) and stained with either CD45-APC/Cy7 (BD Biosciences) or CD45.1-AF700 (Biolegend) and CD45.2-FITC (ThermosFisher) and CD4- APC, CD8-BV570, CD25-BV421, cKit-BV785 (BioLegend) and CD44-PE/Cy7 (ThermoFisher). Flow cytometric analysis was performed on a BD FACS Aria III (BD Biosciences).

### Transplantation of bone marrow, lymphoid progenitors and ThyMCs

8 weeks old C57Bl/6 mice received a single dose of 9.5 Gray 12-24 hours prior to the transplantation. For lymphoid progenitor transplantations bone marrow from C57BL/6- Tg(UBC-GFP)30Scha/J donors was lineage depleted (Miltenyi) following the manufacturer’s instructions. The cells were subsequently stained with biotinylated lineage antibodies (CD3e, B220, CD4, CD8a, Gr-1, Cd11b), cKit-APC and CD135-BV421 (all from BD Bioscience) for 30 minutes at 4°C. This was followed by a 15-minute incubation with Streptavidin-PE/Cy7 (Biolegend). Lineage- CD135+ cKit+GFP+ lymphoid progenitors were sorted on a BD FACS Aria III and 40 000 cells were injected into each lethally irradiated recipient along with 10^6^ nucleated whole bone marrow cells from B6.SJL-Ptprca Pepcb/BoyJ donors. In the case of the adoptive transfer of ThyMCs 2000-10 000 ThyMCs were injected intrathymically along with a retro-orbital injection of 10^6^ nucleated whole bone marrow cells from B6.SJL-Ptprca Pepcb/BoyJ mice 12 hours post-irradiation. The sham injection consisted of sterile PBS in the same volume as the cell grafts.

### sJTREC analysis

TREC quantification was performed as described previously ^61^. In brief, tissue was harvested and DNA isolated using either homogenization with in a Bullet Blender Storm BBX24 instrument (Next Advance, Inc.) followed by genomic DNA extraction using TRIZOL or using a DNeasy Blood and Tissue kit (Qiagen). 1 μg of DNA per sample was used as input for real-time PCR. A standard curve of mouse sjTREC plasmid was used to calculate the absolute number of single joint TRECS (sjTRECs) per sample. All values were normalized to tissue weight.

### T cell receptor sequencing analysis

CD3+ T cells were sorted from spleens of bone marrow transplant recipients 4 weeks post-transplant. RNA was subsequently extracted using RNeasy isolation kit (Qiagen) and RNA content was quantified using a NanoDrop (ThermoFisher) spectrophotometer. Equal amounts of RNA from each sample was submitted to iRepertoire for sequencing and bioinformatics analysis. In brief, samples were reverse transcribed and amplified using a primer set which specifically amplifies beta TCR RNA. The results generated a range of total reads and numbers of unique CDR3s for each sample.

### Reverse-transcription quantitative PCR analysis

To assess the mRNA expression level of *Ccl19* and *Postn* RT-qPCR analysis was performed. 5000-10 000 ThyMCs were sorted into lysis buffer and the RNA was extracted using the NucleoSpin® RNA XS RNA isolation kit (Macherey-Nagel). Total RNA was reverse transcribed into cDNA using the SuperScript IV First-Strand Synthesis System (Thermo Fisher). qPCR analysis was carried out using iTaq Universal SYBR Green Supermix with primers specific for *Ccl19* (Forward-CTGCCTCAGATTATCTGCCAT, Reverse- AGGTAGCGGAAGGCTTTCAC), *Postn* (Forward-CACGGCATGGTTATTCCTTCA, Reverse-TCAGGACACGGTCAATGACAT) and *Gapdh* (Forward- TGTGTCCGTCGTGGATCTGA, Reverse- TTGCTGTTGAAGTCGCAGGAG). Threshold values (C_T_) were estimated using CFX Maestro 2.3 (Biorad) and transcript levels were normalized by subtracting the corresponding *Gapdh* values. The relative amount of RNA was presented as 2^−ΔΔCt^.

### Culture of bone marrow and thymic mesenchymal cells

Bone marrow and thymic mesenchymal cells were grown under hypoxic conditions (2% O_2_) in α-MEM (ThermoFisher) supplemented with 20% FBS (Gibco), and 1% Penicillin/Streptomycin (Gibco). To assess the colony forming ability of individual ThyMCs, Itgb5+ CD99l2+ and Itgb5- CD99l2- cells were sorted as single cells into wells of a 96-well plate. Their colony forming ability was assessed after 7 days of culture. CFU-F potential was determined by seeding 100 mesenchymal cells in wells of 6-well plates. The number of colonies was counted 6 days post-plating.

### Enzyme-linked immunoassay

The protein levels of Flt3l and Ccl19 were determined by enzyme-linked immunoassays (ELISA) (both from Abcam). ∼10mg of thymic tissue was finely minced in 100 μl of ice cold PBS. Tissue remnants were removed by centrifugation for 10 minutes at 1500 *g* and supernatant was promptly removed for Flt3l and Ccl19 measurement. The ELISAs were carried out in accordance with the instructions provided by the manufacturer. The data was eventually normalized to the amount of tissue collected from each sample.

### Gene editing and lentiviral overexpression

SgRNAs directed towards *GFP* and *Ccl19* were designed using the Broad Institute sgRNA tool (https://portals.broadinstitute.org/gpp/public/analysis-tools/sgrna-design) and cloned into a guide-only lentiviral vector^60^. Thymic CD248- ThyMCs from Cas9-GFP expressing mice were isolated as described above and subsequently cultured under hypoxic conditions (2% O_2_) in α-MEM (ThermoFisher) supplemented with 10% FBS (Gibco), and 1% Penicillin/Streptomycin (Gibco) over night. 12 hours post-plating the cells the cells were infected with the lentiviral sgRNA vectors targeting *GFP* and *Ccl19* respectively. Following another 30 hours in culture the gene edited CD248- ThyMCs were intrathymically injected into HSCT recipients.

Bone marrow stromal cells were isolated from Cas9-GFP mice as previously described and infected with pRecieverLv230_mCherry control or pRecieverLv230_mCherry_Ccl19 vectors (Genecopeia, EX-NEG-Lv230 and EX-Mm06316- Lv230) after the second passage in culture. The cells were subsequently grown in the presence of 5μg/ml Puromycin (Gibco) for a minimum of 96 hours to ensure that only transformed cells were present in the cultures before being used for intrathymic injections. 50 000-150 000 *mCherry* or *Ccl19 mCherry* bone marrow stromal cells were subsequently intrathymically injected into either HSCT recipients or 2 year old mice.

### Transwell migration assay

6.0 × 10^4^ *Ccl19 mCherry* and *mCherry* overexpressing bone marrow stromal cells were seeded in 48-well transwell plates (Corning, pore size 5μm). For Cas9-GFP CD248- ThyMCs experiments, 5000 cells were seeded in 96-well transwell plates (Corning, pore size 5μm). Knockout of *GFP* and *Ccl19* was achieved through infection with lentiviral sgRNA vectors as described above. 24 hours later thymocytes from UBC-GFP mice were stained with CD45.2-APC (BioLegend) and CCR7-PE (BD Biosciences). CD45+CCR7+ and CD45+CCR7- cells were sorted and 5.0 × 10^5^ (48-well format) or 2.5 × 10^5^ (96-well format) of each cell type was placed in the upper chamber of the transwell plates. 18 hours later the number of GFP+ cells in the upper and lower chamber was assessed by flow cytometry and CountBright Absolute Counting Beads (ThermoFisher) in accordance with the manufacturer’s instructions.

### Vaccination response

44 or 56 days following HSCT, recipient mice were immunized with 100 μg ovalbumin (OVA, InvivoGen), 100 μg CpG-ODN (Integrated DNA Technologies) and 1 μg GM-CSF (Peprotech). The animals were then challenged 10 days later. 12 days post-vaccination, splenocytes were isolated from these mice by mechanical disruption.

For assessment of the OVA specific CD8+ T cell response, splenocytes were stained with SIINFEKL-APC tetramer (MBL International) together with the following antibodies; CD8-PE (Clone KT15, MBL International) CD4-BV421, Cd11b-PE/Cy7, Gr1-PE/Cy7, B220-PE/Cy7 (all from BioLegend), CD3e-BUV737 and CD45-APC/Cy7 (BD Biosciences). Animals were considered as responders if their CD8+ T_CTL_ population contained higher than background proportions of SINFEKL+ cells. Background level SINFEKL labeling was established by analysis of 3 non-vaccinated animals of the same age group. Flow cytometric analysis was performed on a BD LSR II (BD Biosciences).

To assess cytokine production in ovalbumin specific T cells, irradiated splenocytes from naive C57Bl/6 mice were pulsed with 0.5mg/ml ovalbumin (InvivoGen). CD8+ T cells were isolated from the spleens of the vaccinated mice using CD8 magnetic microbeads (Miltenyi) in accordance with the manufacturer’s instructions. Eventually 200 000 CD8+ T cells from vaccinated animals were seeded with 50 000 ovalbumin pulsed splenocytes on IFNγ ELISpot assay plates (BD Biosciences). The cells were incubated for 48 hours and the plates were developed as described by the manufacturer.

### Adult thymic organ culture

Thymic lobes from adult mice were dissected, cut into smaller pieces and grown on sponge supported (Surgifoam gelatin sponges, Mckesson) 0.8μm filters (Isopore membrane filters, MilliporeSigma) at the air-liquid interface in RPMI-1640 (ThermoFisher) supplemented with 10% FBS (Gibco), 50μM β-mercaptoethanol (MilliporeSigma), 10mM HEPES (Gibco), 1% Penicillin/Streptomycin (Gibco) and 1% Non-essential amino acids (Gibco) as described previously ^62^. In order to induce CreER expression, tissue from *Postn-CreER-tdT/iDTA* mice was grown in the presence of 4-HT (MilliporeSigma) at a final concentration of 4μM for 8 days. The 4-HT containing medium was replaced every 48 hours. Depletion efficiency was determined by flow cytometry after 8 days of culture. Penk+ ThyMC were deleted *in vitro* by growing tissue from *Penk-Cre-tdT/iDTR* animals in the presence of 100ng/ml diphtheria toxin (MilliporeSigma) for 8 days ^75^. Medium containing diphtheria toxin was refreshed every 48 hours. Flow cytometry was used to assess level of depletion on day 8. Evaluation of thymic stroma and hematopoietic components was done through flow cytometric confocal image analysis day 8 of culture.

### Kidney capsule transplantation

After 8 days of adult thymic organ culture in the presence of 4-hydroxytamoxifen, individual pieces of thymic tissue were implanted under the kidney capsule of 8 weeks old syngeneic B6.Cg-Foxn1nu/J mice, as described previously ^20^.

### Thymus sectioning, immunostaining and confocal imaging

Thymic tissue preparation, immunostaining and volumetric confocal tissue imaging was performed as recently described ^63^. Briefly, thymi were fixed in 4% methanol-free formaldehyde (Thermo) for 4 hours (4°C, rotation), PBS washed and embedded in 4% low-melting agarose (Sigma). 200μm thick sections were blocked/permeabilized for 2 hours at room temperature (TBS containing 20% DMSO, 0.05% Tween-20 and 0.5% IGEPAL CA- 630, 10% donkey serum), immunostained overnight using primary [anti-keratin 5 (chicken, Biolegend, 905901), anti-Pdgfra (goat, R&D, AF1062), anti-RFP (rabbit, Rockland, 600-401- 379), anti-GFP (chicken, AVESm GFP-1020)], secondary antibodies (488, 555, 633, 680 Biotium) and DAPI (D1306, Thermo Fischer Scientific) and then were optically cleared (2,2- thiodiethanol, Sigma) to enable deep-tissue 3D imaging. Confocal tissue imaging was performed on a Leica SP8 equipped with two photomultiplier tubes, 3 HyD detectors and 3 laser lines (405, argon, white-light laser) with a 20x multiple-immersion objective (NA 0.75, FWD 0.680 mm). Images were acquired at 400Hz, 8-bit with 1024×1024 resolution.

### Statistical analysis

Statistical tests and figures were generated using Prism v9.1.0. Sample sizes were chosen based on previous experiments and no statistical methods were employed to predetermine sample size. Statistical tests used to compare gene expression differences in the scRNAseq data set is specified in its section of the methods. Fraction of cell populations from scRNA-Seq analysis was compared with beta regression using betareg R package [doi: 10.18637/jss.v034.i02], multiple testing correction of regression coefficients’ p-values was performed using Benjamini-Hochberg adjustment. For pairwise comparisons statistical significance was calculated using unpaired, 2-tailed t-test and for comparisons between three or more groups one-way ANOVA followed by Dunnet’s or Tukey’s post-hoc analysis. All data is presented as mean ±SEM and statistical significance was set to p<0.05 and significance is denoted as follows: *<0.05, **<0.01, ***<0.001 and ****<0.0001.

## Data availability

scRNAseq data generated for this work can be made available upon request.

## Code availability

All code for the scRNA-Seq analysis can be made available upon request.

## Acknowledgments

The authors thank Oscar J Benavidez from the Division of Pediatric-Congenital Cardiology at Massachusetts General Hospital for providing human thymic tissue. We were supported with expert technical assistance by the HSCI-CRM Flow Cytometry facility and the CRM Multiphoton Microscopy Core at Massachusetts General Hospital and the Bauer Core Facility at Harvard University. We thank Youmna Kfoury for sharing her Cas9-GFP bone marrow stroma. We acknowledge Stephen Voinea for providing assistance with coding. We thank Igor Adameyko for assistance in identifying neural crest progenitors in the dataset. K. G and N.B were supported by the Swedish Research Council. N.S was a recipient of the AACR- Millennium Fellowship in Prostate Cancer Research.

## Contributions

K.G, N.B and D.T.S conceived the study. K.G designed and performed most of the experiments with technical assistance and discussions with N.B, N.S, E.W.S and H.G. I.S and P.K did the scRNA-seq analysis. T.Z performed the kidney capsule grafts. K.A.K designed and performed the CRISPR knockout studies. K.D.K developed and performed the 3D tissue- wide confocal thymus imaging. K.G, I.S, N.B, P.K, C.P.L and D.T.S interpreted the data and wrote the manuscript. All authors read, edited and approved the manuscript.

## Supplemental fig legends

<u>Supplemental fig 1.</u>

(A) Gating strategy for flow cytometric sorting of human thymus stromal cells (CD45- CD235-Lineage-CD4-CD8-). Percentages refer to percent of parent gate.

(B) Gating strategy for flow cytometric sorting of murine thymus stromal cells (CD45- Ter119-). Percentages refer to percent of parent gate.

(C) Distribution of number of expressed genes and number of UMIs per cell in the human and mouse data sets.

(D) UMAP embedding of human thymus cell populations including *CD3E+ CD4+ CD8B+ PTPRC+* hematopoietic cells with detailed annotation. (n=3, total number of cells=21034) Three independent experiments.

(E) Expression of marker genes for human hematopoietic cells visualized on joint embedding. All *CD3E+ CD4+ CD8B+ PTPRC+* cells were excluded from future analysis.

(F) UMAP embedding of murine thymus cell populations including *Cd3e+ Cd4+ Cd8a+ Ptprc+* hematopoietic cells with detailed annotation. (n=4, total number of cells=13952) Four independent experiments.

(G) Expression of marker genes for murine hematopoietic cells visualized on joint embedding. All *Cd3e+ Cd4+ Cd8a+ Ptprc+* cells were excluded from future analysis.

<u>Supplemental fig 2.</u>

(A) The top most differentially expressed genes in each major human stromal cell subset shown as a dot plot.

(B) Expression of marker genes for each major human stromal cell subset shown as a dot plot.

(C) The top most differentially expressed genes in each major murine stromal cell subset shown as a dot plot.

(D) Expression of marker genes for each major murine stromal cell subset shown as a dot plot.

(E) Genes defining neural crest-like (NC) cells in murine tissue shown as a dot plot.

(F) Gating strategy for flow cytometric validation of human and mouse thymus stromal cell proportions. The label above the flow plots refer to the previous gates and the percentages are of parent gate.

<u>Supplemental fig 3.</u>

(A) Dot plots displaying genes differentially expressed across human and murine thymic ThyMC subsets.

(B) Expression of marker genes *CD248, PENK, COL15A1, POSTN* that define each ThyMC subtype in human (left) and mouse (right). Presented as UMAP embeddings and bar graphs displaying expression in each cell type. Statistical significance was assessed using the DESeq2 with Benjamini-Hochberg FDR control of these comparisons. (human n= 3, murine n= 4).

(C) UMAP representation of conos joint embedding showing overlap between our human samples (colored dots) and a publicly available human dataset (gray dots). ThyMCs colored by origin (left). Overlays of individual population markers (right) indicate transcriptionally similar populations between the two sample sets.

(D) UMAP embedding showing an overlay thymic ThyMCs from a publically available murine data set with the presently analyzed samples, indicating three transcriptionally comparable subsets are present in both data sets.

(E) UMAPs showing clustering of ThyMCs from a publically available data set in to three subsets expressing the pre-selected marker genes *Cd28, Penk*, and *Postn*.

(F) Normalized enrichment score for GSEA pathways specific to murine Postn+ ThyMC presented as UMAP.

(G) Expression of *Ccl19, Flt3l*, and *Il15* across ThyMC subsets and other stromal cell types. (n= 4).

<u>Supplemental fig 4.</u>

(A) UMAP embedding of murine thymus cell populations including *Cd3e+ Cd4+ Cd8b+ Ptprc+* hematopoietic cells with detailed annotation (left). ThyMC marker genes *Cd99l2, Itgb5, Pdgfra* and *Cd248* presented as UMAP graphs (right). (n=4) Four independent experiments.

(B) FACS plot validating the presence of both CD99l2 and Itgb5 at the protein level on non-hematopoietic, non-epithelial, non-endothelial thymic cells.

(C) Flow cytometric plots demonstrating the overlap between CD99l2 and Itgb5 and other cell type defining markers. Labels to the right of the flow plots refer to parent gate and the percentages are of the parent gate.

(D) Image of fixed and Giemsa stained CD45-Ter119-CD31- EpCam- CD99l2+ Itgb5+ ThyMCs grown in aMEM with 20% FBS under hypoxic conditions for 6 days (left).

(E) Bar graphs displaying colony forming ability of individually sorted CD45-Ter119-CD31- EpCam- CD99l2+ Itgb5+ and CD45-Ter119-CD31- EpCam- CD99l2- Itgb5- cells grown of 7 days in aMEM with 20% FBS under hypoxic conditions.

(F) Bar graph representation of bone marrow and thymic ThyMCs fibroblast colony forming units (CFU-F) after 6 days of hypoxic culture in aMEM with 20% FBS (n=6 per group).

(G) Flow cytometric analysis confirmation at the protein level of CD248 and Pdgfra.

(H) Validation of increased *Postn* expression by qPCR on sorted CD248+ ThyMC compared to CD248- ThyMC. Statistical significance was calculated by unpaired student’s t-test. n=4

<u>Supplemental fig 5.</u>

(A) Gating strategy for FACS analysis of murine thymus tdTomato+ (Penk+) ThyMCs in *Penk-Cre-tdT* mice. Percentages refer to percent of parent gate.

(B) FACS plots showing overlap between tdTomato+ cells and thymic epithelium (EpCam), endothelium (CD31), hematopoietic cells (CD45), and myeloid cells (F4/80, CD11b) in *Penk-Cre-tdT* mice.

(C) Representative histograms from flow cytometric analysis of intracellular Cre protein staining in hematopoietic cells (CD45) and ThyMC (CD248-).

(D) Bar graphs showing *Postn* and *Ccl19* expression as determined by qPCR on sorted tdTomato+ (Penk ThyMC) and tdTomato- (Postn+ ThyMC) cells from *Penk-Cre-tdT* mice.

(E) Gating strategy for flow cytometric analysis of murine thymus tdTomato+ (Postn+) ThyMCs after *in vivo* administration of 4-hydroxytamoxifen in *Postn-CreERT-tdT* mice. Percentages refer to percent of parent gate.

(F) Flow cytometric analysis of overlap between tdTomato+ cells and thymic epithelium (EpCam), endothelium (CD31), and hematopoietic cells (CD45) in *Postn-CreER-tdT* mice.

(G) Bar graphs showing *Postn* and *Ccl19* expression as determined by qPCR on sorted tdTomato+ (Postn+ ThyMC) and tdTomato- (Penk+ ThyMC) cells from 4- hydroxytamoxifen induced *Postn-CreERT-tdT* mice.

(H) Detailed ThyMC annotation presented as ThyMC specific UMAP embedding (left). UMAP showing expression of marker genes *DPP4 (Dpp4), PDPN (Pdpn), LTBR (Ltbr), S100A4 (S100a4)* in human (top) and mouse (bottom).

(I) Representative histogram showing staining of S100A4 and Ltbr on thymic ThyMCs from *Penk-Cre-tdT* mice.

(J) Bar graphs showing mean fluorescent intensity (MFI) on CD248+ ThyMC, Penk+ ThyMC and Postn+ ThyMC as determined by flow cytometry in *Penk-Cre-tdT* mice.

(K) FACS plots showing staining for DPP4 and Pdpn on CD248+ ThyMC, Penk+ ThyMC and Postn+ ThyMC from *Penk-Cre-tdT* mice.

(L) Bar graphs showing the percentage of phenotypic cortical fibroblasts (cFib; DPP4+ Pdpn+) and medullary fibroblasts (mFib; Dpp4- Pdpn+) within CD248+ ThyMC, Penk+ ThyMC and Postn+ ThyMC subsets from *Penk-Cre-tdT* mice. Statistical significance is based on beta regression with Benjamini-Hochberg FDR control of these comparisons.

<u>Supplemental fig 6</u>

(A) UMAP embedding showing stromal cell (*Cd3e-, Cd4-, Cd8a-, Ptprc-)* compositional differences between Control (n=4, total number of stromal cells=5451), Transplantation (n=3, total number of stromal cells=8057), IL7RKO (n=4, total number of stromal cells=4551) and Aging (n=4, total number of stromal cells=16178) samples.

(B) Bar graphs comparing proportional shifts of different stromal cell subsets between Control, Transplantation, IL7RKO and Aging samples as determined by scRNAseq. (Control n= 4, Transplantation n= 3, IL7RKO n=4, Aging n=4). Statistical significance is based on beta regression with Benjamini-Hochberg FDR control of these comparisons.

(C) Dot plot showing expression of neural crest-like (NC) cell marker genes (rows) across different stress states (columns). Total number of NCs per condition noted in parenthesis.

(D) Flow cytometry plots showing the presence of cKit+ stromal cells within the CD45- Ter119-CD31-Epcam- population in Control and IL7R KO thymic tissue.

(E) Expression of marker genes defining murine osteogenic smooth muscle cells (oSMC) presented as dot plot.

(F) Bar graphs showing the difference in proportions within the MC population of Penk+ ThyMCs in Control, Transplantation, and Aging samples as determined by scRNAseq. (Control n= 4, Transplantation n= 3, IL7RKO n=4, Aging n=4). Statistical significance is based on beta regression with Benjamini-Hochberg FDR control of these comparisons.

(G) Volumetric confocal microscopy images from Day 3 post-HSCT Postn-CreER-tdTomato thymus stained with DAPI (white; cell nuclei), and tdTomato (red; Postn+ ThyMCs).

(H) Bar graphs showing *Postn* and *Ccl19* expression as determined by qPCR on sorted CD248- ThyMC 2 months and 22-27 months old mice (n=4). Statistical significance was calculated by unpaired two-tailed student’s t-test.

(I) Bar graph comparing proportional shifts of the total ThyMC population between Control, Transplantation, IL7RKO and Aging samples as determined by scRNAseq. (Control n= 4, Transplantation n= 3, IL7RKO n=4, Aging n=4) Statistical significance is based on beta regression with Benjamini-Hochberg FDR control of these comparisons.

(J) Heatmap displaying gene set enrichment analysis (GESA) of differentially expressed genes between Transplantation (n=3) and Control (n=4) samples, as well as Aging (n=4) and Control (n=4) samples across different murine ThyMC subtypes. Significance level marked as dot corresponds to FDR between 0.05 and 0.25 following official GSEA User Guide.

(K) Bar graphs showing the results of ELISA measurements of Ccl19 and Flt3l protein levels in whole thymic tissue in Control (n=5) and Transplantation (n=5) samples. Statistical significance was calculated by unpaired two-tailed student’s t-test.

<u>Supplemental fig 7</u>

(A) Representative flow plots showing the extent of tdTomato depletion ThyMC in *Postn-CreERT-tdT/iDTA* and *Postn-CreERT-tdT* samples respectively after 4-HT induction. The analyzed cells are CD248- ThyMC, gated on as described in Supplemental fig 5E. Percentages refer to percent of parent gate.

(B) Bar graphs showing the flow cytometric quantification of stromal cell subsets in Control (iDTA Ctrl n=4) and Postn-CreER-tdT/iDTA (Postn iDTA n=6) mice 6 days post-bone marrow transplantation. Statistical significance was calculated by unpaired two-tailed student’s t-test. Two independent experiments.

(C) FACS analysis of thymocyte subsets Control (iDTA Ctrl n=4) and Postn-CreER-tdT/iDTA (Postn iDTA n=6) mice 6 days post-bone marrow transplantation. Statistical significance was calculated by unpaired two-tailed student’s t-test. Two independent experiments.

(D) Gating strategy for flow cytometric analysis of *in vitro* depletion of Postn+ ThyMC in murine adult thymic organ cultures (ATOC) using tissue from *Postn-CreERT- tdT/iDTA* mice. Percentages refer to percent of parent gate.

(E) Volumetric confocal microscopy images from a Postn-CreER-tdTomato day 8 ATOC stained with DAPI (white; cell nuclei), Keratin-5 (green; mTECs), Keratin-8 (cyan; cTEC), and tdTomato (red; Postn+ ThyMCs).

(F) Bar graphs showing FACS analysis of T cell developmental stages in *iDTA* Control and *Postn-CreERT-iDTA* samples after 8 days of adult thymic organ culture (ATOC) in the presence of 4-hydroxytamoxifen. (*iDTA* Control= 6, *Postn-CreERT-iDTA*= 6) Statistical significance was calculated by unpaired two-tailed student’s t-test. Two independent experiments.

(G) Flow cytometric analysis of thymic stromal cells subsets in *iDTA* Control and *Postn- CreERT-iDTA* samples after 8 days of adult thymic organ culture (ATOC) in the presence of 4-hydroxytamoxifen presented as bar graphs. (*iDTA* Control= 6, *Postn- CreERT-iDTA*= 6) Two independent experiments. Statistical significance was calculated by unpaired two-tailed student’s t-test.

<u>Supplemental fig 8</u>

(A) Schematic illustration of experimental design for kidney capsule graft of adult thymic organ cultures (ATOCs) into athymic nude mice.

(B) Flow cytometric of assessment of T cells in peripheral blood of treatment Naive, and nude mouse recipients of iDTA Ctrl or Postn iDTA kidney capsule ATOC grafts 8 weeks post-implantation. Statistical significance was calculated using Student’s t-test. (n= 4 Naive, n=4 iDTA Ctrl and n=3 Postn Ctrl).

(C) Quantification of *de novo* generated T cells in kidney capsule ATOC grafts from nude mouse recipients 10 weeks following implantation quantified using signal-joint T cell receptor excision circles (sjTRECs). Statistical significance was determined by Mann-Whitney U test.

(D) Representative flow plots showing the percent of tdTomato labeled MC in *Penk-Cre- tdT/iDTR* and *Penk-Cre-tdT* mice respectively after diphtheria toxin mediated depletion. The analyzed cells are CD248- MC, gated on as described in Supplemental fig 5A. Percentages refer to percent of parent gate.

(E) Bar graphs showing FACS quantification of stromal cell subsets in Control (iDTR Ctrl n=7) and Penk-Cre-tdT/iDTR (Penk iDTR n=7) mice 6 days post-bone marrow transplantation and diphtheria toxin injection. Statistical significance was calculated by unpaired two-tailed student’s t-test. Two independent experiments.

(F) Flow cytometric analysis of thymocyte subsets Control (iDTR Ctrl n=7) and Penk-Cre- tdT/iDTR (Postn iDTA n=7) mice 6 days post-bone marrow transplantation and diphtheria toxin injection. Statistical significance was calculated by unpaired two-tailed student’s t-test. Two independent experiments.

(G) Flow cytometric analysis of thymic stromal cells subsets in *iDTR* Control and *Penk-Cre- tdT/iDTR* samples after 8 days of ATOC in the presence of diphtheria toxin presented as bar graphs. (*iDTR* Control= 3, *Penk-Cre-tdT/iDTR*= 3). Statistical significance was calculated by unpaired two-tailed student’s t-test.

(H) Bar graphs showing FACS analysis of T cell developmental stages in *iDTR* Control and *Penk-Cre-tdT/iDTR* samples after 8 days of ATOC in the presence of diphtheria toxin. (*iDTR* Control= 3, *Penk-Cre-tdT/iDTR*= 3). Statistical significance was determined by Student’s t-test.

<u>Supplemental fig 9</u>

(A) Representative flow plots showing the presence of GFP+ Penk+ ThyMCs and tdTomato+ Postn+ ThyMCs in the thymi of bone marrow transplantation recipients 6 days post- intrathymic injection of 5 000 cells.

(B) FACS quantification of thymus endothelium, epithelium and ThyMCs 6 days after intrathymic injection of Sham, GFP+ (Penk+) ThyMC or tdTomato+ (Postn+) ThyMCs displayed as bar graphs. 4000-8000) (Sham=9, Penk+ ThyMC=14, Postn+ ThyMCs=10) Three independent experiments. Statistical significance was determined by one-way ANOVA followed by Tukey’s post-hoc analysis.

(C) Flow cytometric plots showing GFP labeled cells 6 days after adoptive transfer of 2000- 4000 CD8+ T cells or CD248- ThyMCs.

(D) Bar graphs displaying the flow cytometric quantification of endothelial cells, epithelial cells and ThyMCs 6 days following an intrathymic sham injection or adoptive transfer of CD8+ T cells and CD248- ThyMCs. (Dose of CD8+ T cells and CD248- ThyMCs: 2000- 4000) (Sham n= 4, CD8+ T cells n=6, CD248- ThyMC n=8) Two independent experiments. Statistical significance was determined by one-way ANOVA followed by Tukey’s post-hoc analysis.

(E) Bar graphs showing the results of the FACS quantification of the proportion of GFP labeled CCR7+ or CCR7- thymocytes that migrated to the bottom of transwell wells after 18 hours in the presence of GFP KO Ctrl or Ccl19 KO Cas9-GFP CD248- ThyMCs. (n= 3).

(F) FACS quantification of early endothelium, epithelium and MCs following an intrathymic injection of Sham, Cas9-GFP CD248- MC GFP knockout control (KO Ctrl) or Cas9-GFP CD248- MC Ccl19 knockout (Ccl19 KO). (Dose of GFP KO Ctrl and Ccl19 KO cells: 4000) Values are presented as percent of Sham treated animals. (n= 15 Sham, n=11 GFP KO Ctrl, n=10 Ccl19 KO). Three independent experiments.

(G) Flow cytometric plots showing mCherry labeled control (mCherry Ctrl) or Ccl19 mCherry overexpressing (Ccl19 OE) bone marrow stromal cells adoptively transferred to the thymus 6 days earlier.

(H) Bar graphs showing the results of the FACS quantification of the proportion of GFP labeled CCR7+ or CCR7- thymocytes that migrated to the bottom of transwell wells after 18 hours in the presence of Ccl19 OE or mCherry Ctrl bone marrow stromal cells. (n= 3).

(I) Bar graphs showing the results of FACS quantification of endothelial cells, epithelial cells and MCs following an intrathymic sham injection or adoptive transfer of Cas9-GFP bone marrow stromal cells following infection with of either mCherry Ctrl or Ccl19 OE vectors. (Dose of mCherry Ctrl and Ccl19 OE cells: 50 000) Values are presented as percent of Sham treated animals.(n= 6 Sham, n=9 mCherry Ctrl, n=9 Ccl19 OE). Two independent experiments.

<u>Supplemental fig 10</u>

(A) Flow cytometric analysis of myeloid and B cells in peripheral blood following adoptive transfer of Sham, CD8+ T cells or CD248- MCs 2-16 weeks post-transplantation. (Dose of CD8+ T cells and CD248- MCs: 10 000) (Sham=7, CD8+ T cells=9, CD248- ThyMCs=9). Two independent experiments. Gray shaded area denotes the peripheral blood parameters of untreated control mice (n=12) housed in the animal facility at the same time as the transplant recipients.

(B) Flow cytometric plots demonstrating presence of CD8+ T cells and CD248- MCs in the thymi of bone marrow transplantation recipients 16 weeks post-intrathymic injection of 10 000 cells.

(C) Bar graphs showing thymocyte cell count in thymi isolated from Sham, CD8+ T cells and CD248- MC treated bone marrow transplant recipients 4 weeks post-transfer. (Dose of CD8+ T cells and CD248- MCs: 10 000) (Sham=8, CD8+ T cells=12, CD248- ThyMCs= 14) Two independent experiments. Statistical significance was determined by one-way ANOVA followed by Tukey’s post-hoc analysis.

(D) Gating strategy for flow cytometric analysis of SINFEKL+ CD8+ T_CTL_ cells following ovalbumin vaccination in recipients of Sham, CD8+ T cell or CD248- MC intrathymic injections. Analyzed cells are gated on singlet lymphocytes and percentages refer to percent in parent gate.

(E) Bar graphs showing number of thymocytes per thymic lobe after an intrathymic injection of bone marrow stromal cells overexpressing either mCherry Ctrl or Ccl19 OE. (Dose of mCherry Ctrl and Ccl19 OE cells: 150 000) Statistical significance was determined using Student’s t-test comparing Aged mCherry Ctrl with Aged Ccl19 OE. (Young n= 9, Aged mCherry Ctrl n=9, Aged mCherry Ccl19 OE n=7). Two independent experiments.

(F) Flow cytometric analysis showing mCherry Ctrl or Ccl19 OE bone marrow stromal cells adoptively transferred to the thymus of 24 months old ovalbumin vaccine recipients 69 days earlier.

(G) FACS quantification of absolute number of SINFEKL+ CD8+ T_CTL_ cells in ovalbumin vaccine responsive animals presented as a bar graph. (Dose of mCherry Ctrl and Ccl19 OE cells: 150 000). (Responding Young n= 9, Responding Aged mCherry Ctrl n=2, Responding Aged mCherry Ccl19 OE n=5. Two independent experiments.

