## Supplemental table 1 for "Thymic mesenchymal niche cells drive T cell immune regeneration"

Table 1 Human samples

| Sample Identifier | Sample Type | Sample Description | Digestion time | Age | Sex | 10X kit |
|---|---|---|---|---|---|---|
| SCG_HTS1_20190722 | Human thymus niche | Sorted (CD45-/Cd235a-/Lin-) human thymus niche cells | 90 minutes | 2.5 months | Female | Version 2 |
| SCG_HTS1_20200302 | Human thymus niche | Sorted (CD45-/Cd235a-/Lin-) human thymus niche cells | 90 minutes | 4 months | Male | Version 2 |
| SCG_HTS1_20200309 | Human thymus niche | Sorted (CD45-/Cd235a-/Lin-) human thymus niche cells | 90 minutes | 2 months | Female | Version 2 |

Table 1 Murine samples

| Sample.ID | Sample.type | Sample.description | Genotype | Irradiation | Transplant | Age | Sex | 10X kit |
|---|---|---|---|---|---|---|---|---|
| SCG_65 | Murine Thymus niche | Steady-state; sorted (CD45-/Ter119-) murine thymus niche from C57BL/6 mouse | WT | NO | NO | 8 weeks | Female | Version 2 |
| SCG_MTH2_C | Murine Thymus niche | Steady-state; sorted (CD45-/Ter119-) murine thymus niche from C57BL/6 mouse | WT | NO | NO | 8 weeks | Female | Version 2 |
| SCG_MTH2_IR | Murine Thymus niche | Sorted (CD45-/Ter119-) murine thymus niche cells from irradiated and transplanted 3 C57BL/6 mice | WT | YES | YES | 8 weeks | Female | Version 2 |
| SCG_MTH3_C | Murine Thymus niche | Steady-state; sorted (CD45-/Ter119-) murine thymus niche from C57BL/6 mouse | WT | NO | NO | 8 weeks | Male | Version 2 |
| SCG_MTH3_IL7_C | Murine Thymus niche | Sorted (CD45-/Ter119-) murine thymus niche cells from irradiated and transplanted 4 IL7RKO mice | IL7 receptor knockout (IL7RKO) | NO | NO | 8 weeks | Female | Version 2 |
| SCG_MTH4_C | Murine Thymus niche | Steady-state; sorted (CD45-/Ter119-) murine thymus niche from C57BL/6 mouse | WT | NO | NO | 8 weeks | Male | Version 2 |
| SCG_MTH5_IL7_C | Murine thymus niche | Sorted (CD45-/Ter119-) murine thymus niche cells from irradiated and transplanted 4 IL7RKO mice | IL7 receptor knockout (IL7RKO) | NO | NO | 8 weeks | Male | Version 2 |
| SCG_MTH5_IR | Murine Thymus niche | Sorted (CD45-/Ter119-) murine thymus niche cells from irradiated and transplanted 3 C57BL/6 mice | WT | YES | YES | 8 weeks | Male | Version 2 |
| SCG_MTH7_IR | Murine Thymus niche | Sorted (CD45-/Ter119-) murine thymus niche cells from irradiated and transplanted 3 C57BL/6 mice | WT | YES | YES | 8 weeks | Female | Version 2 |
| SCG_MTH8_IL7_C1 | Murine Thymus niche | Sorted (CD45-/Ter119-) murine thymus niche cells from irradiated and transplanted 4 IL7RKO mice | IL7 receptor knockout (IL7RKO) | NO | NO | 8 weeks | Female | Version 2 |
| SCG_MTH8_IL7_C2 | Murine Thymus niche | Sorted (CD45-/Ter119-) murine thymus niche cells from irradiated and transplanted 4 IL7RKO mice | IL7 receptor knockout (IL7RKO) | NO | NO | 8 weeks | Male | Version 2 |
| SCG_MTH9_Old_M1 | Murine Thymus niche | Steady-state; sorted (CD45-/Ter119-) murine thymus niche from 2 C57BL/6 mice | WT | NO | NO | 96 weeks | Male | Version 3 |
| SCG_MTH9_Old_M2 | Murine Thymus niche | Steady-state; sorted (CD45-/Ter119-) murine thymus niche from 2 C57BL/6 mice | WT | NO | NO | 96 weeks | Male | Version 3 |
| SCG_MTH9_Old_F3 | Murine Thymus niche | Steady-state; sorted (CD45-/Ter119-) murine thymus niche from 2 C57BL/6 mice | WT | NO | NO | 96 weeks | Female | Version 3 |
| SCG_MTH9_Old_F4 | Murine Thymus niche | Steady-state; sorted (CD45-/Ter119-) murine thymus niche from 2 C57BL/6 mice | WT | NO | NO | 96 weeks | Female | Version 3 |
