## Supplemental table 2 for "Thymic mesenchymal niche cells drive T cell immune regeneration"

### Figure 4B

No intrathymically injcted cells detcted by flow

**Experiment 1**

| Mouse ID | ETP % of Sham | Endothelial % of Sham | Epithelialthelial % of Sham | ThyMC % of Sham |
|---|---|---|---|---|
| 1 Sham | 91.82 | 15.41 | 118.06 | 129.68 |
| 2 Sham | 120.32 | 26.37 | 70.77 | 125.23 |
| 5 Sham | 63.86 | 236.03 | 93.80 | 80.63 |
| 10 Sham | 123.99 | 122.20 | 117.37 | 64.46 |
| 3 Penk ThyMC | 139.54 | 205.86 | 114.08 | 85.03 |
| 6 Penk ThyMC | 264.81 | 123.67 | 153.86 | 239.41 |
| 7 Penk ThyMC | 129.57 | 84.78 | 536.01 | 89.68 |
| 11 Penk ThyMC | 89.37 | 62.94 | 129.81 | 87.14 |
| 8 Postn ThyMC | 521.82 | 238.29 | 157.49 | 244.21 |
| 9 Postn ThyMC | 376.23 | 250.35 | 72.14 | 105.65 |

**Experiment 2**

| Mouse ID | ETP % of Sham | Endothelial % of Sham | Epithelialthelial % of Sham | ThyMC % of Sham |
|---|---|---|---|---|
| 1 Sham | 29.95 | 42.85 | 76.78 | 58.08 |
| 4 Sham | 141.64 | 105.17 | 110.97 | 74.67 |
| 13 Sham | 128.41 | 151.98 | 112.25 | 167.25 |
| 2 Penk ThyMC | 67.71 | 139.91 | 115.19 | 64.15 |
| 5 Penk ThyMC | 182.99 | 88.32 | 57.76 | 145.37 |
| 8 Penk ThyMC | 215.96 | 292.87 | 128.17 | 71.48 |
| 10 Penk ThyMC | 158.57 | 110.34 | 103.20 | 132.20 |
| 11 Penk ThyMC | 109.39 | 70.13 | 62.02 | 143.93 |
| 3 Postn ThyMC | 502.07 | 462.71 | 185.46 | 272.79 |
| 6 Postn ThyMC | 275.50 | 284.60 | 58.80 | 93.26 |
| 7 Postn ThyMC | 95.24 | 97.58 | 142.56 | 99.93 |
| 9 Postn ThyMC | 200.20 | 261.70 | 108.48 | 43.13 |
| 12 Postn ThyMC | 145.10 | 124.69 | 102.92 | 34.25 |

**Experiment 3**

| Mouse ID | ETP% of Sham | Endothelial cells % of Sl | Epithelial cells % of sham | ThyMCs % of Sham |
|---|---|---|---|---|
| 4 Sham | 95.41 | 85.73 | 85.55 | 119.04 |
| 5 Sham | 104.59 | 114.27 | 114.45 | 80.96 |
| 1 Penk ThyMC | 158.79 | 131.69 | 101.00 | 182.52 |
| 2 Penk ThyMC | 241.44 | 125.03 | 189.85 | 150.10 |
| 7 penk ThyMC | 234.56 | 254.60 | 243.77 | 120.06 |
| 10 Penk ThyMC | 138.72 | 169.79 | 82.50 | 33.02 |
| 11 Penk ThyMC | 122.18 | 176.75 | 97.62 | 81.72 |
| 3 Postn ThyMC | 492.16 | 560.21 | 301.86 | 167.94 |
| 6 Postn ThyMC | 175.40 | 335.98 | 300.60 | 100.06 |
| 12 Postn ThyMC | 84.55 | 160.22 | 83.23 | 47.24 |

**Experiment 1 Absolute numbers**

| Mouse ID | ETP | Endothelial | Epithelial | ThyMCs |
|---|---|---|---|---|
| 1 Sham | 16.80 | 115.27 | 811.15 | 8111.48 |
| 2 Sham | 22.01 | 197.18 | 486.21 | 7833.31 |
| 5 Sham | 11.68 | 1765.19 | 644.43 | 5043.40 |
| 10 Sham | 22.69 | 913.88 | 806.36 | 4031.82 |
| 3 Penk ThyMC | 25.53 | 1539.61 | 783.80 | 5318.65 |
| 6 Penk ThyMC | 48.45 | 924.92 | 1057.06 | 14974.96 |
| 7 Penk ThyMC | 23.71 | 634.09 | 3682.59 | 5609.24 |
| 11 Penk ThyMC | 16.35 | 470.71 | 891.87 | 5450.30 |
| 8 Postn ThyMC | 95.47 | 1782.144 | 1082.02 | 15275.52 |
| 9 Postn ThyMC | 68.84 | 1872.312 | 495.61 | 6608.16 |

**Experiment 2 Absolute numbers**

| Mouse ID | ETP | Endothelial | Epithelial | ThyMCs |
|---|---|---|---|---|
| 1 Sham | 61.57 | 200.64 | 2498.88 | 5107.20 |
| 4 Sham | 291.20 | 492.48 | 3611.52 | 6566.40 |
| 13 Sham | 264.00 | 711.63 | 3653.03 | 14707.02 |
| 2 Penk ThyMC | 139.20 | 655.13 | 3748.79 | 5641.38 |
| 5 Penk ThyMC | 376.20 | 413.58 | 1879.90 | 12783.32 |

| | | | | |
|---|---|---|---|---|
| 8 Penk ThyMC | 444.00 | 1371.36 | 4171.22 | 6285.40 |
| 10 Penk ThyMC | 326.00 | 516.69 | 3358.47 | 11625.48 |
| 11 Penk ThyMC | 224.90 | 328.40 | 2018.27 | 12656.96 |
| 3 Postn ThyMC | 1032.20 | 2166.64 | 6035.64 | 23987.80 |
| 6 Postn ThyMC | 566.40 | 1332.63 | 1913.52 | 8200.80 |
| 7 Postn ThyMC | 1958.00 | 456.92 | 4639.54 | 8787.00 |
| 9 Postn ThyMC | 411.60 | 1225.39 | 3530.30 | 3792.88 |
| 12 Postn ThyMC | 298.30 | 583.87 | 3349.57 | 3011.54 |

**Experiment 3 Absolute numbers**

| Mouse ID | ETP | Endothelial | Epithelial | ThyMCs |
|---|---|---|---|---|
| 4 Sham | 233.02 | 143.08 | 1022.00 | 8993.60 |
| 5 Sham | 255.46 | 190.69 | 1367.24 | 6116.60 |
| 1 Penk ThyMC | 387.83 | 219.77 | 1206.58 | 13789.44 |
| 2 Penk ThyMC | 589.68 | 208.66 | 2268.00 | 11340.00 |
| 7 penk ThyMC | 572.88 | 424.89 | 2912.14 | 9070.60 |
| 10 Penk ThyMC | 338.80 | 283.36 | 985.60 | 2494.80 |
| 11 Penk ThyMC | 298.41 | 294.98 | 1166.20 | 6174.00 |
| 3 Postn ThyMC | 1202.04 | 934.92 | 3606.12 | 12688.20 |
| 6 Postn ThyMC | 428.40 | 560.7 | 3591.00 | 7560.00 |
| 12 Postn ThyMC | 206.50 | 272.7858 | 994.27 | 3569.16 |

### Figure 4C

No intrathymically injected cells detected by flow

**Experiment 1 Percent of Sham**

| Mouse ID | ETP % of Sham | ThyMC % of Sham | Endothelial % of Sham | Epithelial % of Sham |
|---|---|---|---|---|
| Sham 3 | 100.00 | 100.00 | 100.00 | 100.00 |
| CD248- ThyMC 4 | 214.05 | 274.10 | 255.33 | 225.78 |
| CD248- ThyMC 6 | 188.25 | 89.21 | 171.32 | 120.57 |

**Experiment 2 Percent of Sham**

| Mouse ID | ETP % of Sham | ThyMC % of Sham | Endothelial % of Sham | Epithelial % of Sham |
|---|---|---|---|---|
| Sham 1 | 77.21 | 83.39 | 77.55 | 77.55 |
| Sham 2 | 97.36 | 99.69 | 124.46 | 124.46 |
| Sham 3 | 125.42 | 116.92 | 97.98 | 97.98 |
| CD8+ T cell 4 | 124.22 | 221.29 | 128.63 | 120.33 |
| CD8+ T cell 5 | 181.17 | 207.49 | 186.63 | 192.06 |
| CD8+ T cell 6 | 132.58 | 322.82 | 188.65 | 153.99 |
| CD8+ T cell 10 | 102.82 | 114.10 | 90.04 | 145.34 |
| CD8+ T cell 11 | 157.97 | 273.34 | 161.60 | 247.38 |
| CD8+ T cell 12 | 46.97 | 178.67 | 107.36 | 50.73 |
| CD248- ThyMC 7 | 218.93 | 530.32 | 508.20 | 192.00 |
| CD248- ThyMC 8 | 303.58 | 335.08 | 375.28 | 302.32 |
| CD248- ThyMC 9 | 225.80 | 573.25 | 494.83 | 437.20 |
| CD248- ThyMC 13 | 108.11 | 88.55 | 146.41 | 119.21 |
| CD248- ThyMC 14 | 238.52 | 165.10 | 262.55 | 88.69 |
| CD248- ThyMC 15 | 119.96 | 90.76 | 142.12 | 94.42 |

**Experiment 1 Absolute numbers**

| Mouse ID | ETP | ThyMC | Endothelial | Epithelial |
|---|---|---|---|---|
| Sham 3 | 134.13 | 16253.40 | 1357.08 | 1783.14 |
| CD248- ThyMC 4 | 252.50 | 14500.00 | 2325.00 | 2150.00 |
| CD248- ThyMC 6 | 287.10 | 44550.00 | 3465.00 | 4026.00 |

**Experiment 1 Absolute numbers**

| Mouse ID | ETP | ThyMC | Endothelial | Epithelial |
|---|---|---|---|---|
| Sham 1 | 82.00 | 2296.00 | 1230.00 | 319.80 |
| Sham 2 | 103.40 | 2744.80 | 1974.00 | 451.20 |
| Sham 3 | 133.20 | 3219.00 | 1598.40 | 754.80 |
| CD8+ T cell 4 | 131.92 | 6092.80 | 2040.00 | 612.00 |
| CD8+ T cell 5 | 192.40 | 5712.80 | 2960.00 | 976.80 |
| CD8+ T cell 6 | 140.80 | 8888.00 | 2992.00 | 783.20 |
| CD8+ T cell 10 | 109.20 | 3141.60 | 1428.00 | 739.20 |
| CD8+ T cell 11 | 167.76 | 7525.90 | 2563.00 | 1258.20 |
| CD8+ T cell 12 | 49.88 | 4919.20 | 1702.80 | 258.00 |
| CD248- ThyMC 7 | 232.50 | 14601.00 | 8060.00 | 976.50 |
| CD248- ThyMC 8 | 322.40 | 9225.60 | 5952.00 | 1537.60 |
| CD248- ThyMC 9 | 239.80 | 15783.20 | 7848.00 | 2223.60 |
| CD248- ThyMC 13 | 114.81 | 2438.10 | 2322.00 | 606.30 |
| CD248- ThyMC 14 | 253.31 | 4545.70 | 4164.00 | 451.10 |
| CD248- ThyMC 15 | 127.40 | 2499.00 | 2254.00 | 480.20 |

### Figure 4D

**Experiment 1**

| Mouse ID | ETP % of Sham | Endothelial % Sham | Epithelial % of Sham | ThyMC % of Sham |
|---|---|---|---|---|
| 1 Sham | 114.40 | 140.36 | 88.06 | 137.00 |
| 2 Sham | 123.81 | 129.07 | 132.37 | 132.00 |
| 7 Sham | 119.79 | 95.10 | 86.70 | 105.00 |
| 8 Sham | 89.85 | 94.06 | 78.96 | 79.10 |
| 14 Sham | 52.16 | 41.41 | 113.92 | 45.90 |
| 3 GFP KO Ctrl | 140.33 | 205.68 | 96.48 | 129.15 |
| 4 GFP KO Ctrl | 111.43 | 160.84 | 103.25 | 117.61 |
| 9 GFP KO Ctrl | 192.13 | 181.59 | 120.51 | 232.00 |
| 10 GFP KO Ctrl | 133.48 | 97.82 | 72.45 | 126.60 |
| 15 GFP KO Ctrl | 104.39 | 102.00 | 128.44 | 83.44 |
| 6 Ccl19 KO | 114.73 | 169.15 | 196.60 | 148.56 |
| 11 Ccl19 KO | 105.25 | 57.20 | 43.80 | 103.59 |
| 12 Ccl19 KO | 105.84 | 115.54 | 136.92 | 178.73 |
| 13 Ccl19 KO | 57.27 | 31.30 | 48.98 | 100.81 |

**Experiment 2**

| Mouse ID | ETP % of Sham | Endothelial % Sham | Epithelial % of Sham | ThyMC % of Sham |
|---|---|---|---|---|
| 1 Sham | 83.88 | 88.79 | 96.72 | 85.11 |
| 4 Sham | 121.25 | 87.28 | 86.35 | 85.40 |
| 7 Sham | 58.85 | 124.98 | 53.21 | 36.57 |
| 10 Sham | 116.28 | 81.69 | 128.07 | 159.49 |
| 12 Sham | 106.48 | 75.13 | 96.40 | 130.06 |
| 14 Sham | 111.32 | 121.51 | 148.16 | 111.73 |
| 15 Sham | 101.94 | 120.61 | 91.10 | 91.65 |
| 2 GFP KO Ctrl | 270.19 | 204.39 | 64.53 | 99.18 |
| 5 GFP KO Ctrl | 129.44 | 118.54 | 60.12 | 56.89 |
| 9 GFP KO Ctrl | 114.00 | 61.02 | 101.56 | 64.98 |
| 13 GFP KO Ctrl | 134.14 | 120.14 | 71.74 | 167.83 |
| 3 KO | 138.50 | 109.50 | 65.86 | 62.43 |
| 6 KO | 90.73 | 84.20 | 126.00 | 167.37 |
| 8 KO | 116.51 | 107.09 | 119.06 | 114.74 |
| 11 KO | 88.21 | 67.75 | 69.06 | 75.86 |

**Experiment 3**

| Mouse ID | ETP % of Sham | Endothelial % Sham | Epithelial % of Sham | ThyMC % of Sham |
|---|---|---|---|---|
| 7 Sham | 125.61 | 98.56 | 101.03 | 96.40 |
| 10 Sham | 137.99 | 101.88 | 88.71 | 177.00 |
| 11 Sham | 69.89 | 78.08 | 97.44 | 76.00 |
| 12 Sham | 66.51 | 121.48 | 112.82 | 51.10 |
| 5 GFP KO Ctrl | 168.96 | 111.79 | 103.46 | 96.87 |
| 8 GFP KO Ctrl | 228.02 | 121.18 | 93.50 | 109.43 |
| 6 Ccl19KO | 126.96 | 130.24 | 132.75 | 103.60 |
| 9 Ccl19KO | 91.73 | 95.48 | 89.06 | 112.23 |

**Experiment 1 Absolute numbers**

| Mouse ID | ETP | Endothelium | Epithelium | ThyMCs |
|---|---|---|---|---|
| 1 Sham | 347.600 | 537.200 | 1137.600 | 18738.800 |
| 2 Sham | 376.200 | 494.000 | 1710.000 | 32908.000 |
| 7 Sham | 364.000 | 364.000 | 1120.000 | 22652.000 |
| 8 Sham | 273.000 | 360.000 | 1020.000 | 17160.000 |
| 14 Sham | 158.480 | 158.480 | 1471.600 | 12791.600 |
| 3 GFP KO Ctrl | 426.400 | 787.200 | 1246.400 | 26928.800 |
| 4 GFP KO Ctrl | 338.580 | 615.600 | 1333.800 | 24521.400 |
| 9 GFP KO Ctrl | 583.800 | 695.000 | 1556.800 | 48372.000 |
| 10 GFP KO Ctrl | 405.600 | 374.400 | 936.000 | 26395.200 |
| 15 GFP KO Ctrl | 317.200 | 390.400 | 1659.200 | 17397.200 |
| 6 Ccl19 KO | 348.600 | 647.400 | 2539.800 | 30975.600 |
| 11 Ccl19 KO | 319.800 | 218.940 | 565.800 | 21598.800 |
| 12 Ccl19 KO | 321.600 | 442.200 | 1768.800 | 37265.400 |
| 13 Ccl19 KO | 174.020 | 119.780 | 632.800 | 21018.000 |

**Experiment 2 Absolute numbers**

| Mouse ID | ETP | Endothelium | Epithelium | ThyMCs |
|---|---|---|---|---|
| 1 Sham | 170.720 | 931.200 | 2095.200 | 39188.000 |
| 4 Sham | 246.760 | 915.400 | 1870.600 | 39322.400 |
| 7 Sham | 119.780 | 1310.800 | 1152.600 | 16837.000 |
| 10 Sham | 236.640 | 856.800 | 2774.400 | 73440.000 |
| 12 Sham | 216.700 | 788.000 | 2088.200 | 59888.000 |
| 14 Sham | 226.560 | 1274.400 | 3209.600 | 51448.000 |
| 15 Sham | 207.460 | 1265.000 | 1973.400 | 42200.400 |

| | | | | |
|---|---|---|---|---|
| 2 GFP KO Ctrl | 545.220 | 2143.600 | 1398.000 | 45668.000 |
| 5 GFP KO Ctrl | 263.440 | 1243.200 | 1302.400 | 26196.000 |
| 9 GFP KO Ctrl | 232.000 | 640.000 | 2200.000 | 29920.000 |
| 13 GFP KO Ctrl | 273.000 | 1260.000 | 1554.000 | 77280.000 |
| 3 KO | 281.880 | 1148.400 | 1426.800 | 28744.800 |
| 6 KO | 184.644 | 883.080 | 2729.520 | 77068.800 |
| 8 KO | 237.120 | 1123.200 | 2579.200 | 52832.000 |
| 11 KO | 179.520 | 710.600 | 1496.000 | 34931.600 |

**Experiment 2 Absolute numbers**

| Mouse ID | ETP | Endothelium | Epithelium | ThyMCs |
|---|---|---|---|---|
| 7 Sham | 113.880 | 1533.000 | 26280.000 | 49494.000 |
| 10 Sham | 125.100 | 1584.600 | 23074.000 | 41700.000 |
| 11 Sham | 63.360 | 1214.400 | 25344.000 | 43507.200 |
| 12 Sham | 60.300 | 1889.400 | 29346.000 | 30431.400 |
| 5 GFP KO Ctrl | 153.180 | 1738.800 | 26910.000 | 39992.400 |
| 8 GFP KO Ctrl | 206.720 | 1884.800 | 24320.000 | 45174.400 |
| 6 Ccl19KO | 115.100 | 2025.760 | 34530.000 | 42771.160 |
| 9 Ccl19KO | 83.160 | 1485.000 | 23166.000 | 46332.000 |

### Figure 4E

**Experiment 1**

No intrathymically injected cells detected by flow

| Mouse ID | | ETP % Sham | Endothelial % Sham | Epithelial % Sham | ThyMC % Sham |
|---|---|---|---|---|---|
| Sham | | | | | |
| | 1 | 127.47 | 187.39 | 121.69 | 103.05 |
| | 4 | 99.52 | 94.36 | 88.83 | 105.90 |
| | 7 | 73.01 | 18.25 | 89.48 | 91.05 |
| mCherry Ctrl | | | | | |
| | 2 | 77.05 | 48.35 | 42.91 | 55.03 |
| | 5 | 123.36 | 87.32 | 86.62 | 93.56 |
| | 8 | 130.16 | 187.29 | 129.28 | 64.02 |
| | 10 | 77.72 | 51.02 | 50.65 | 52.90 |
| | 12 | 116.72 | 171.58 | 115.08 | 115.23 |
| Ccl19 OE | | | | | |
| | 3 | 122.90 | 131.85 | 105.06 | 92.05 |
| | 6 | 181.93 | 62.15 | 101.86 | 92.39 |
| | 9 | 154.64 | 102.80 | 50.98 | 40.48 |
| | 11 | 179.03 | 154.77 | 144.37 | 88.49 |

**Experiment 2**

| Mouse ID | | ETP % Sham | Endothelial % Sham | Epithelial % Sham | ThyMC % Sham |
|---|---|---|---|---|---|
| Sham | | | | | |
| | 1 | 125.29 | 126.40 | 136.98 | 106.50 |
| | 6 | 59.84 | 62.83 | 51.34 | 59.10 |
| | 11 | 114.87 | 110.77 | 111.68 | 134.40 |
| mCherry Ctrl | | | | | |
| | 3 | 133.64 | 168.53 | 153.22 | 125.35 |
| | 8 | 90.37 | 53.63 | 106.10 | 141.52 |
| | 10 | 87.80 | 69.47 | 75.59 | 115.70 |
| | 13 | 21.36 | 21.79 | 9.80 | 58.41 |
| | 15 | 69.28 | 27.41 | 61.68 | 98.78 |
| Ccl19 OE | | | | | |
| | 2 | 117.09 | 99.70 | 91.89 | 91.25 |
| | 7 | 123.65 | 160.51 | 137.22 | 185.13 |
| | 9 | 195.57 | 121.59 | 158.54 | 121.19 |
| | 12 | 132.52 | 122.33 | 139.37 | 145.72 |
| | 14 | 97.25 | 54.82 | 59.23 | 67.27 |

**Experiment 1 Absolute numbers**

| Mouse ID | | ETP | Endothelium | Epithelium | ThyMCs |
|---|---|---|---|---|---|
| Sham | | | | | |
| | 1 | 116.20 | 10209.00 | 15189.00 | 86320.00 |
| | 4 | 90.72 | 5140.80 | 11088.00 | 88704.00 |
| | 7 | 66.56 | 994.50 | 11169.00 | 76270.50 |
| mCherry Ctrl | | | | | |
| | 2 | 70.24 | 2634.00 | 5355.80 | 46095.00 |
| | 5 | 112.45 | 4757.50 | 10812.50 | 78369.00 |
| | 8 | 118.65 | 10203.90 | 16136.40 | 53629.80 |
| | 10 | 70.85 | 2779.50 | 6322.00 | 44308.50 |
| | 12 | 106.40 | 9348.00 | 14364.00 | 96520.00 |
| Ccl19 OE | | | | | |
| | 3 | 112.03 | 7183.10 | 13114.10 | 77103.00 |
| | 6 | 165.84 | 3385.90 | 12714.40 | 77392.00 |
| | 9 | 140.97 | 5600.70 | 6362.70 | 33909.00 |
| | 11 | 163.20 | 8432.00 | 18020.00 | 74120.00 |

**Experiment 2 Absolute numbers**

| Mouse ID | | | | |
|---|---|---|---|---|
| Sham | ETP | Endothelium | Epithelium | ThyMCs |
| 1 | 822.00 | 1397.40 | 6123.90 | 28030.20 |
| 6 | 392.60 | 694.60 | 2295.20 | 15553.00 |
| 11 | 753.60 | 1224.60 | 4992.60 | 35372.10 |
| mCherry Ctrl | | | | |
| 3 | 876.80 | 1863.20 | 6850.00 | 32989.60 |
| 8 | 592.90 | 592.90 | 4743.20 | 37244.90 |
| 10 | 576.00 | 768.00 | 3379.20 | 30451.20 |
| 13 | 140.16 | 240.90 | 438.00 | 15373.80 |
| 15 | 454.50 | 303.00 | 2757.30 | 25997.40 |
| Ccl19 OE | | | | |
| 2 | 768.20 | 1102.20 | 4108.20 | 24014.60 |
| 7 | 811.20 | 1774.50 | 6134.70 | 48722.70 |
| 9 | 1283.10 | 1344.20 | 7087.60 | 31894.20 |
| 12 | 869.40 | 1352.40 | 6230.70 | 38350.20 |
| 14 | 638.00 | 606.10 | 2647.70 | 17704.50 |

### Figure 4F

**T helper cells/ul blood**

| Weeks | Sham | | | | | | | CD8+ T cells | | | | | | | | | CD248- MC | | | | | | | | |
|---|---|---|---|---|---|---|---|---|---|---|---|---|---|---|---|---|---|---|---|---|---|---|---|---|---|
| 2 | 2.62 | 0.50 | 2.93 | 0.91 | 1.00 | 2.93 | 0.51 | 3.30 | 0.75 | 0.10 | 2.86 | 2.59 | 0.90 | 0.74 | 1.90 | 1.00 | 2.25 | 2.59 | 0.69 | 2.13 | 1.56 | 0.30 | 2.58 | 6.40 | 1.64 |
| 4 | 27.15 | 15.46 | 20.71 | 2.41 | 2.70 | 36.48 | 5.25 | 49.91 | 36.32 | 5.62 | 11.40 | 0.93 | 8.83 | 3.74 | 17.57 | 27.57 | 99.08 | 46.76 | 137.55 | 59.54 | 53.19 | 82.21 | 59.26 | 89.25 | 42.85 |
| 8 | 294.34 | 290.57 | 491.07 | 357.68 | 773.70 | 482.16 | 570.57 | 630.66 | 454.02 | 698.04 | 494.17 | 496.40 | 449.36 | 206.59 | 483.75 | 672.75 | 940.48 | 840.88 | 1528.71 | 1482.62 | 881.79 | 927.63 | 1191.87 | 594.92 | 529.36 |
| 12 | 736.19 | 704.72 | 560.70 | 617.88 | 336.16 | 643.56 | 480.60 | 873.70 | 731.20 | 790.13 | 679.36 | 649.67 | 591.73 | 683.89 | 845.69 | 936.31 | 1607.42 | 1365.51 | 1210.30 | 1941.28 | 1747.07 | 1339.94 | 1365.32 | 925.73 | 1039.76 |
| 16 | 1621.68 | 980.27 | 1812.29 | 1646.88 | 1163.90 | 1941.04 | 1794.44 | 1341.60 | 1393.84 | 1372.68 | 1859.40 | 1297.80 | 1095.19 | 1103.48 | 1683.93 | 1547.19 | 2333.81 | 2419.22 | 1918.00 | 2220.77 | 2193.87 | 1824.73 | 2203.07 | 1684.79 | 1238.46 |

**T CTL cells/ul blood**

| Weeks | Sham | | | | | | | CD8+ T cells | | | | | | | | | CD248- MC | | | | | | | | |
|---|---|---|---|---|---|---|---|---|---|---|---|---|---|---|---|---|---|---|---|---|---|---|---|---|---|
| 2 | 2.46 | 2.80 | 4.80 | 5.40 | 3.42 | 1.76 | 2.58 | 4.06 | 2.16 | 0.40 | 3.02 | 3.03 | 1.99 | 3.07 | 8.92 | 1.54 | 6.76 | 4.42 | 7.10 | 5.75 | 4.56 | 13.09 | 0.82 | 4.74 | 2.53 |
| 4 | 29.37 | 48.05 | 27.03 | 17.08 | 33.06 | 44.46 | 16.70 | 54.68 | 39.47 | 50.01 | 17.56 | 7.44 | 18.28 | 10.38 | 29.89 | 18.72 | 75.03 | 57.66 | 82.43 | 72.50 | 32.40 | 68.38 | 47.48 | 47.08 | 63.29 |
| 8 | 360.45 | 307.71 | 336.36 | 319.60 | 457.69 | 432.80 | 659.95 | 624.45 | 445.05 | 547.63 | 413.63 | 458.32 | 393.95 | 365.88 | 582.57 | 472.83 | 611.42 | 604.05 | 963.96 | 690.23 | 673.46 | 889.26 | 793.44 | 491.80 | 392.86 |
| 12 | 645.19 | 616.90 | 455.68 | 519.51 | 359.84 | 626.11 | 530.44 | 763.71 | 424.00 | 586.95 | 487.13 | 536.71 | 709.64 | 708.58 | 835.74 | 672.57 | 851.30 | 979.26 | 951.47 | 1174.91 | 1255.35 | 1781.09 | 1182.85 | 894.93 | 642.64 |
| 16 | 1265.66 | 726.36 | 857.18 | 792.04 | 852.88 | 770.64 | 688.10 | 536.64 | 674.91 | 664.73 | 871.09 | 1285.44 | 1373.68 | 947.47 | 1062.47 | 862.62 | 970.46 | 1188.07 | 1728.00 | 1026.09 | 1468.09 | 1948.58 | 1592.41 | 1102.92 | 997.91 |

**Myeloid cells/ul blood**

| Weeks | Sham | | | | | | | CD8+ T cells | | | | | | | | | CD248- MC | | | | | | | | |
|---|---|---|---|---|---|---|---|---|---|---|---|---|---|---|---|---|---|---|---|---|---|---|---|---|---|
| 2 | 542.84 | 324.50 | 998.01 | 530.40 | 171.18 | 544.73 | 382.50 | 857.25 | 364.72 | 396.90 | 346.08 | 471.35 | 288.64 | 629.30 | 246.84 | 582.51 | 560.56 | 579.84 | 640.64 | 471.44 | 267.60 | 291.38 | 394.87 | 337.85 | 594.83 |
| 4 | 2069.25 | 1172.64 | 1443.16 | 1640.31 | 2061.90 | 1242.60 | 1119.36 | 1831.33 | 2070.64 | 1053.36 | 2374.68 | 463.14 | 1792.94 | 3274.84 | 1395.80 | 1705.40 | 1357.02 | 2283.62 | 2226.00 | 1543.05 | 1746.90 | 2904.92 | 1982.84 | 2586.93 | 1182.74 |
| 8 | 2620.71 | 2983.57 | 2203.36 | 2556.80 | 2032.80 | 2026.22 | 2688.40 | 2040.10 | 2511.60 | 4080.21 | 3171.42 | 2094.40 | 3937.03 | 4001.50 | 3752.34 | 2643.28 | 3029.94 | 2190.89 | 2795.91 | 4858.92 | 3575.61 | 3029.94 | 2190.89 | 2795.91 | 4858.92 |
| 12 | 3230.50 | 3270.12 | 2447.50 | 2365.83 | 2269.08 | 1479.76 | 2162.70 | 3361.73 | 3592.00 | 4018.35 | 2570.76 | 1157.72 | 4721.81 | 3495.94 | 2647.29 | 3738.63 | 3663.84 | 5546.24 | 2523.27 | 3843.80 | 1816.49 | 3919.96 | 3190.47 | 3003.97 | 4904.82 |
| 16 | 3280.64 | 3021.16 | 1668.42 | 2803.20 | 1707.55 | 2447.20 | 3632.36 | 3213.60 | 1944.04 | 2758.07 | 4210.77 | 2002.32 | 3936.57 | 3298.69 | 2855.49 | 1953.59 | 2955.04 | 3844.24 | 2460.00 | 3209.58 | 2581.02 | 4013.83 | 3928.41 | 3828.03 | 3917.65 |

**B cells/ul blood**

| Weeks | Sham | | | | | | | CD8+ T cells | | | | | | | | | CD248- MC | | | | | | | | |
|---|---|---|---|---|---|---|---|---|---|---|---|---|---|---|---|---|---|---|---|---|---|---|---|---|---|
| 2 | 189.42 | 119.00 | 80.38 | 26.46 | 28.35 | 36.60 | 22.28 | 327.66 | 57.81 | 5.04 | 14.83 | 26.90 | 21.33 | 1.49 | 32.53 | 347.49 | 311.64 | 290.88 | 178.36 | 137.74 | 46.00 | 314.72 | 151.18 | 32.91 | 103.48 |
| 4 | 1717.70 | 1559.04 | 534.10 | 213.09 | 388.60 | 1991.20 | 198.75 | 1350.56 | 460.46 | 83.60 | 418.88 | 228.78 | 25.27 | 37.47 | 428.59 | 936.49 | 1357.02 | 1421.02 | 2220.75 | 1915.65 | 580.50 | 2164.39 | 1892.54 | 220.09 | 1735.83 |
| 8 | 4517.38 | 4144.28 | 3135.10 | 3461.20 | 3505.04 | 2760.94 | 3246.10 | 4517.38 | 4144.28 | 1995.63 | 4295.68 | 3712.80 | 3029.97 | 3942.58 | 4724.57 | 3296.35 | 4724.10 | 4763.19 | 5591.82 | 3861.00 | 3750.03 | 3254.32 | 4027.36 | 5392.73 | 3823.85 |
| 12 | 4149.60 | 2527.56 | 4387.70 | 4333.29 | 4255.48 | 3992.56 | 4708.10 | 3556.87 | 3080.00 | 3440.43 | 3836.84 | 3180.92 | 4208.26 | 4602.93 | 5193.72 | 4375.83 | 2801.76 | 4060.64 | 2465.43 | 6784.05 | 3980.08 | 8549.53 | 2391.49 | 6478.27 | 5949.03 |
| 16 | 7474.64 | 6331.58 | 4195.23 | 3624.45 | 3802.76 | 7022.40 | 5589.23 | 5475.60 | 3108.63 | 3279.18 | 6033.99 | 4115.88 | 4593.03 | 4439.58 | 6388.36 | 4824.56 | 4852.31 | 4672.74 | 6520.00 | 4636.06 | 3739.17 | 6927.94 | 3130.93 | 7419.73 | 4721.38 |

No intrathymically injected cells detected by flow

### Figure 4G

No intrathymically injected cells detected by flow

**Experiment 1**

| Mouse ID | sjTREC/ mg tissue |
|---|---|
| Sham 1 | 100000 |
| 4 | 80000 |
| 7 | 163000 |
| 11 | 58000 |
| CD8+ T cells 2 | 190000 |
| 5 | 87000 |
| 8 | 90000 |
| 12 | 55000 |
| 14 | 190000 |
| 15 | 87000 |
| CD248- MC 3 | 180000 |
| 6 | 287000 |
| 9 | 295000 |
| 13 | 190000 |
| 15 | 178000 |
| 16 | 180900 |
| 17 | 255000 |
| 18 | 170200 |

**Experiment 2**

| Mouse ID | sjTREC/ mg tissue |
|---|---|
| Sham 1 | 98000 |
| 2 | 150000 |
| 3 | 100500 |
| 4 | 95000 |
| CD8+ T cells 5 | 120000 |
| 6 | 48000 |
| 7 | 110000 |
| 8 | 69000 |
| 9 | 170000 |
| 10 | 58000 |

| | |
|---|---|
| CD248- MC 11 | 215000 |
| 12 | 223000 |
| 13 | 210000 |
| 14 | 130300 |
| 15 | 120000 |
| 16 | 220000 |

### Figure 4 J-K

No intrathymically injected cells detected by flow

**Experriment 1**

| Mouse ID | Tetramer+ CD8+ T cell # |
|---|---|
| Sham 1 | 1832.00 |
| Sham 2 | 2678.40 |
| Sham 3 | 5820.00 |
| Sham 4 | 8280.00 |
| Sham 5 | 6219.84 |
| CD8+ T cell 10 | 19583.92 |
| CD8+ T cell 11 | 17624.20 |
| CD8+ T cell 12 | 7176.68 |
| CD8+ T cell 13 | 15760.00 |
| CD8+ T cell 14 | 1600.00 |
| CD248- ThyMC 6 | 16568.00 |
| CD248- ThyMC 7 | 29400.00 |
| CD248- ThyMC 8 | 25664.00 |
| CD248- ThyMC 9 | 25147.00 |

**Experriment 2**

| Mouse ID | Tetramer+ CD8+ T cell # | IFNg spots per 100 000 seeded CTL |
|---|---|---|
| Sham 1 | 15346.00 | 16.50 |
| 4 | 17853.00 | 23.10 |
| 7 | 12678.00 | 12.10 |
| 8 | 13456.00 | 14.30 |
| CD8+ T cells 2 | 5679.00 | 14.30 |
| 5 | 1045.00 | 19.80 |
| 9 | 4230.00 | 8.80 |
| 11 | 14538.00 | 24.20 |
| 14 | 8945.00 | 15.40 |
| 15 | 13785.00 | 25.30 |
| CD248- ThyMC 3 | 21033.00 | 31.90 |
| 6 | 22440.00 | 34.10 |
| 10 | 15954.00 | 39.60 |
| 12 | 14137.00 | 38.50 |

s

### Figure 4M-O

**Experiment 1**

| Mouse ID | Tetramer+ CD8+ T cell # | ETP # | Epithelial # |
|---|---|---|---|
| Young 1 | 23579.10 | 1785.60 | 5039.00 |
| Young 2 | 28951.02 | 2163.10 | 5917.97 |
| Young 3 | 36922.98 | 2051.34 | 3004.00 |
| Young 4 | 3232.08 | 1533.00 | 3956.50 |
| Young 5 | 12681.90 | 2259.98 | 4618.22 |
| Young 6 | 16628.88 | sample lost | sample lost |
| mCherry Ctrl 4 | 18316.20 | 108.64 | 174.46 |
| mCherry Ctrl 5 | 0.00 | 137.79 | 266.00 |
| mCherry Ctrl 9 | 0.00 | 57.73 | 325.14 |
| mCherry Ctrl 11 | 0.00 | 34.28 | 132.48 |
| Ccl19 OE 1 | 5074.50 | 122.98 | 139.36 |
| Ccl19 OE 3 | 0.00 | 68.54 | 462.40 |
| Ccl19 OE 7 | 3638.04 | 210.00 | 1680.00 |
| Ccl19 OE 10 | 3113.28 | 198.48 | 975.86 |

**Experiment 2**

| Mouse ID | Tetramer+ CD8+ T cell # | ETP # | Epithelial # |
|---|---|---|---|
| Young 1 | 4359.78 | 1976.58 | 10008.00 |
| Young 2 | 14760.00 | 3644.20 | 7315.00 |
| Young 3 | 4447.68 | 2553.80 | 13560.00 |
| mCherry 1 | 0.00 | 242.60 | 419.30 |
| mCherry 4 | 8886.24 | 180.81 | 385.63 |
| mCherry 7 | 0.00 | 235.74 | 501.81 |
| mCherry 9 | 0.00 | 104.66 | 913.50 |
| mCherry 11 | 0.00 | 174.46 | 536.80 |
| Ccl19 OE 2 | 35285.00 | 888.80 | 1373.60 |
| Ccl19 OE 6 | 9516.00 | 543.60 | 1268.40 |
| Ccl19 OE 10 | 0.00 | 593.40 | 1573.80 |
