## Supplemental figures 1-10 for "Thymic mesenchymal niche cells drive T cell immune regeneration"

### Supplemental Fig 1

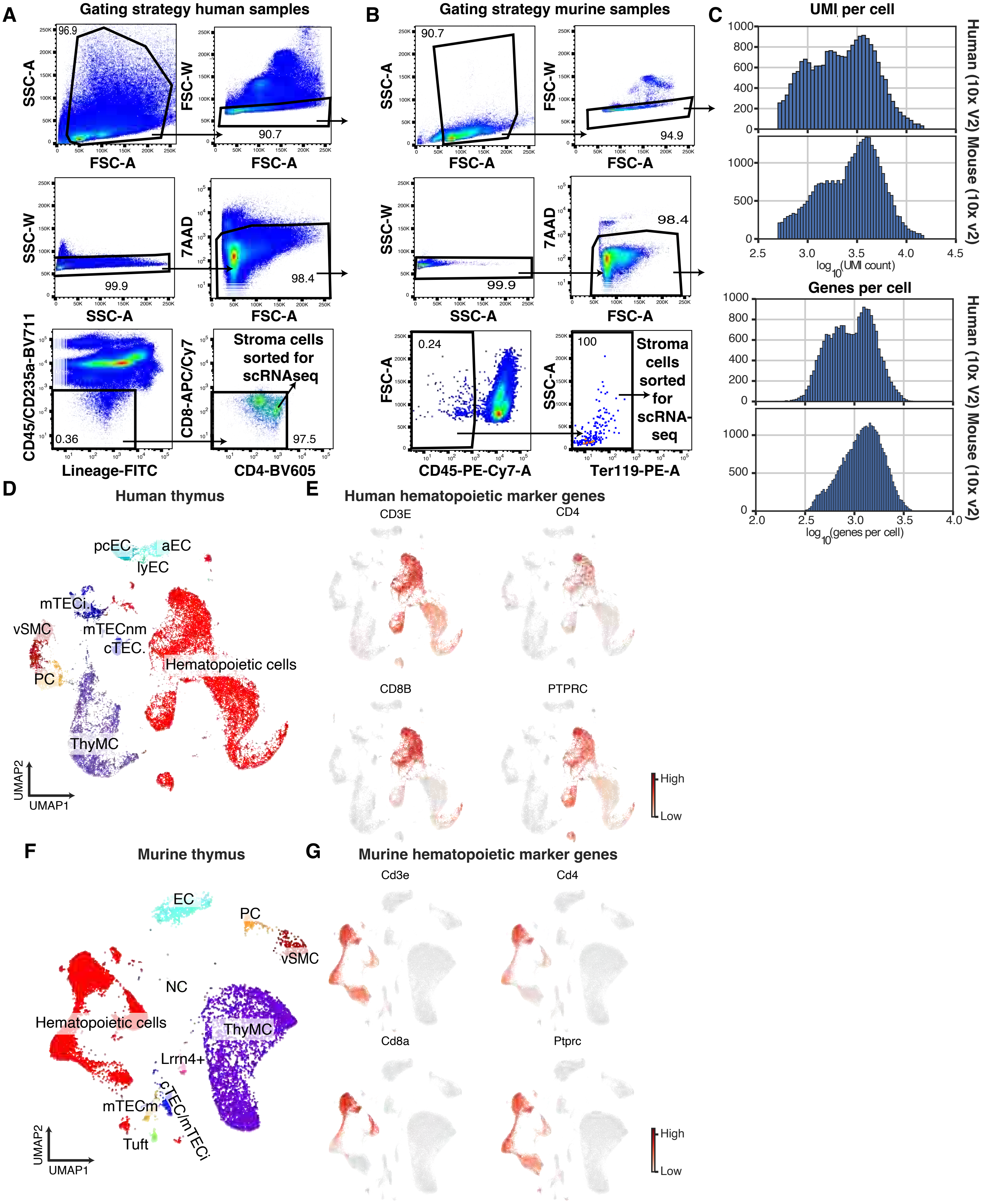

#### Supplemental Fig 2

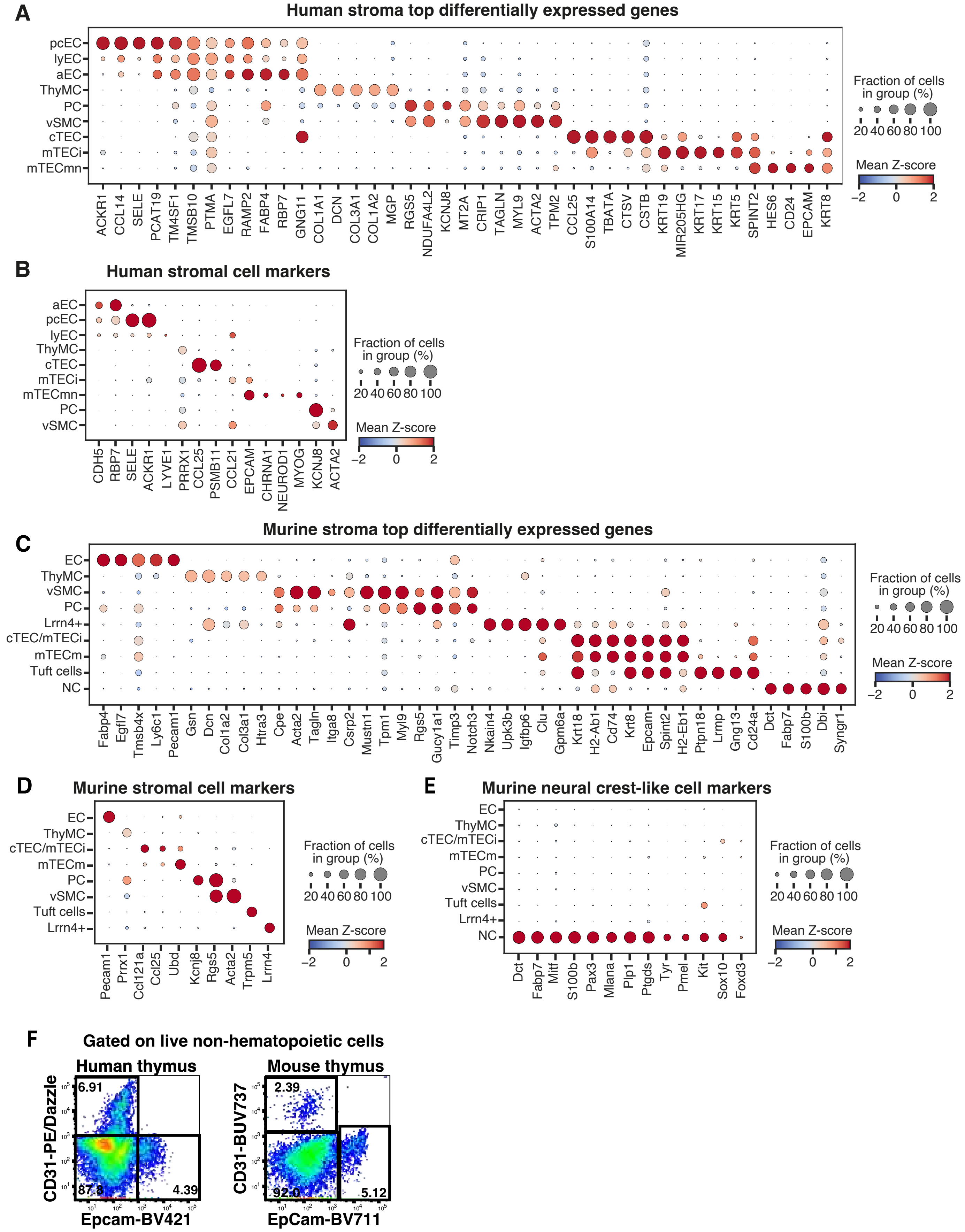

Supplemental Fig 3

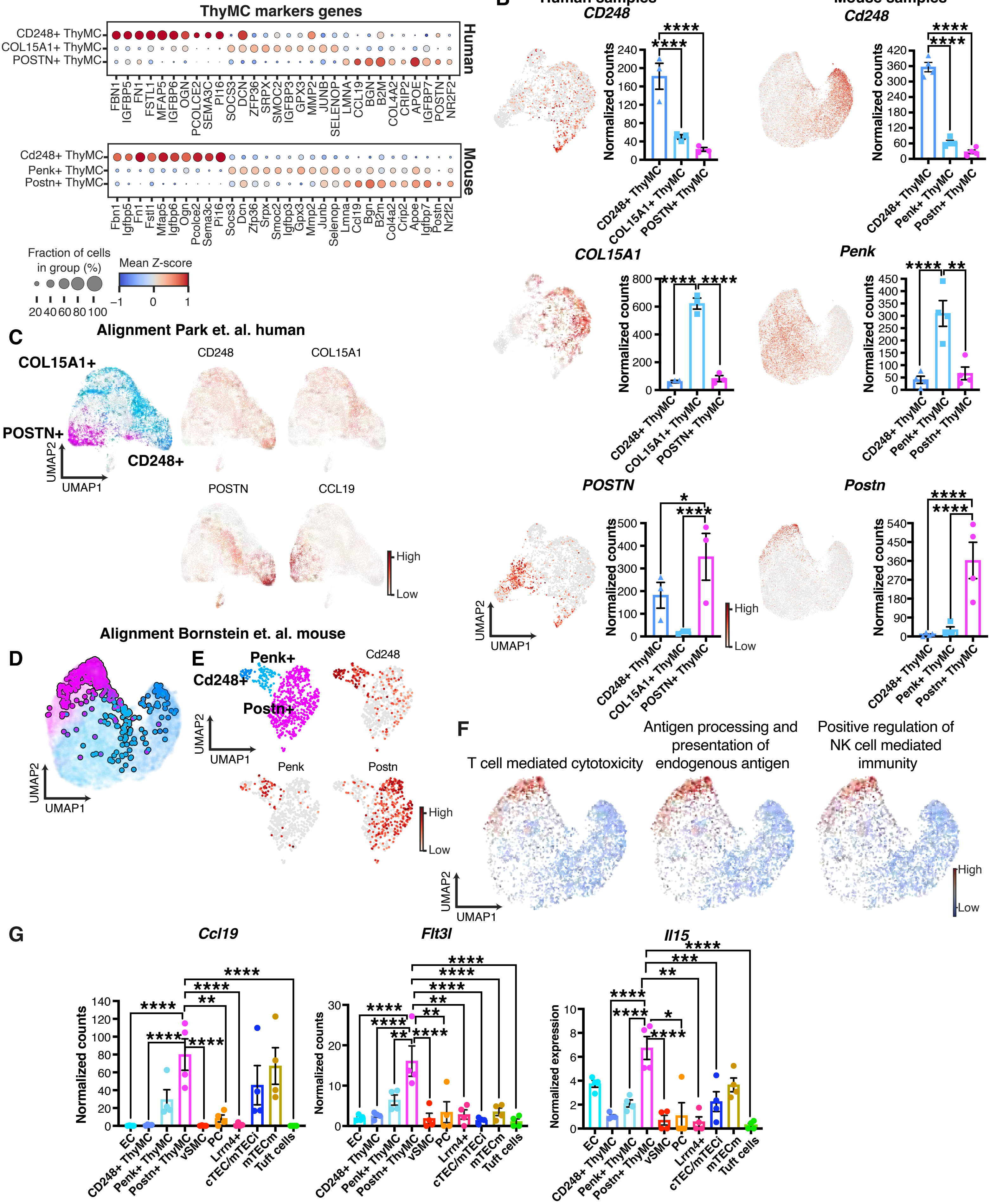

#### Supplemental Fig 4

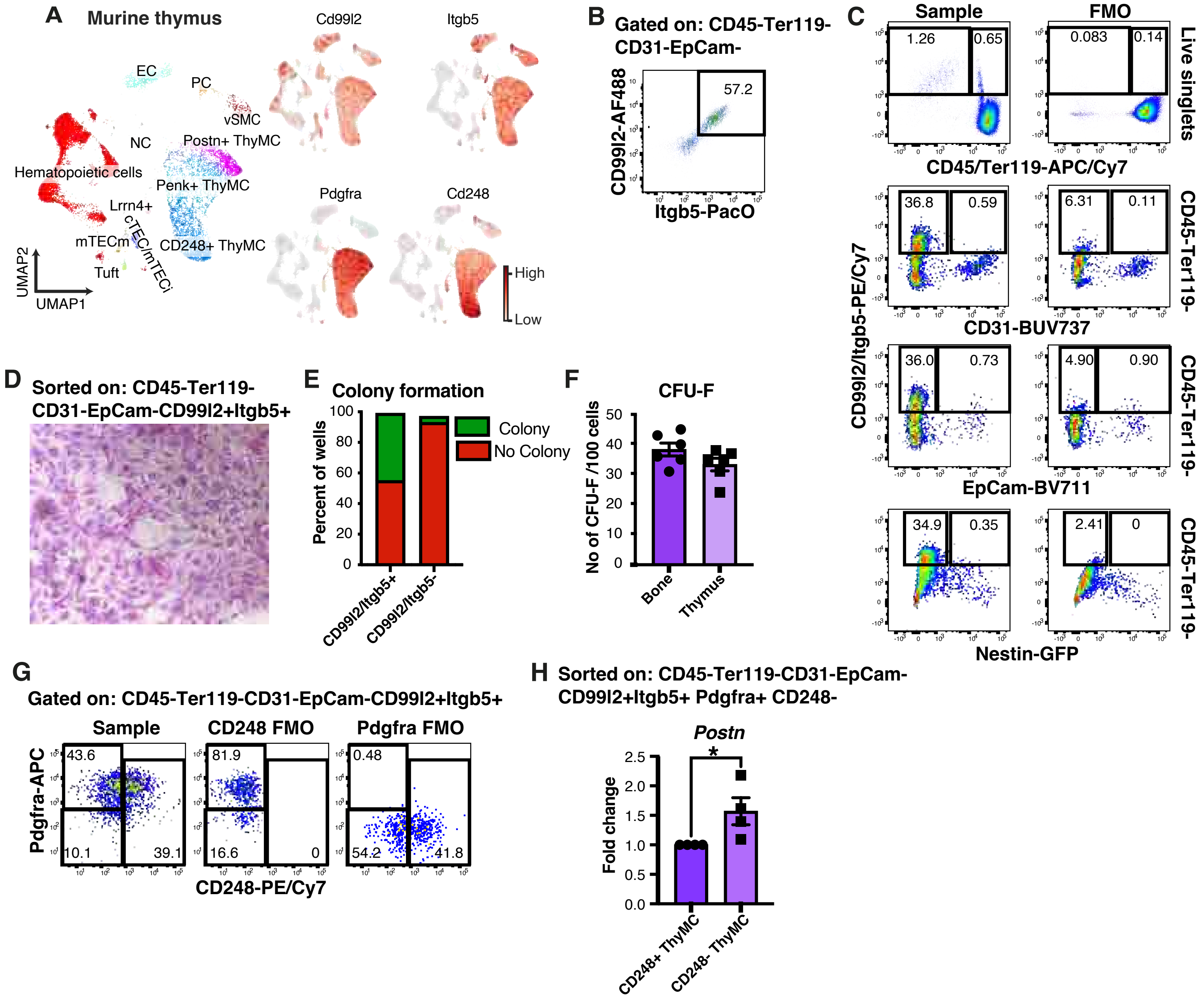

#### Supplemental Fig 5

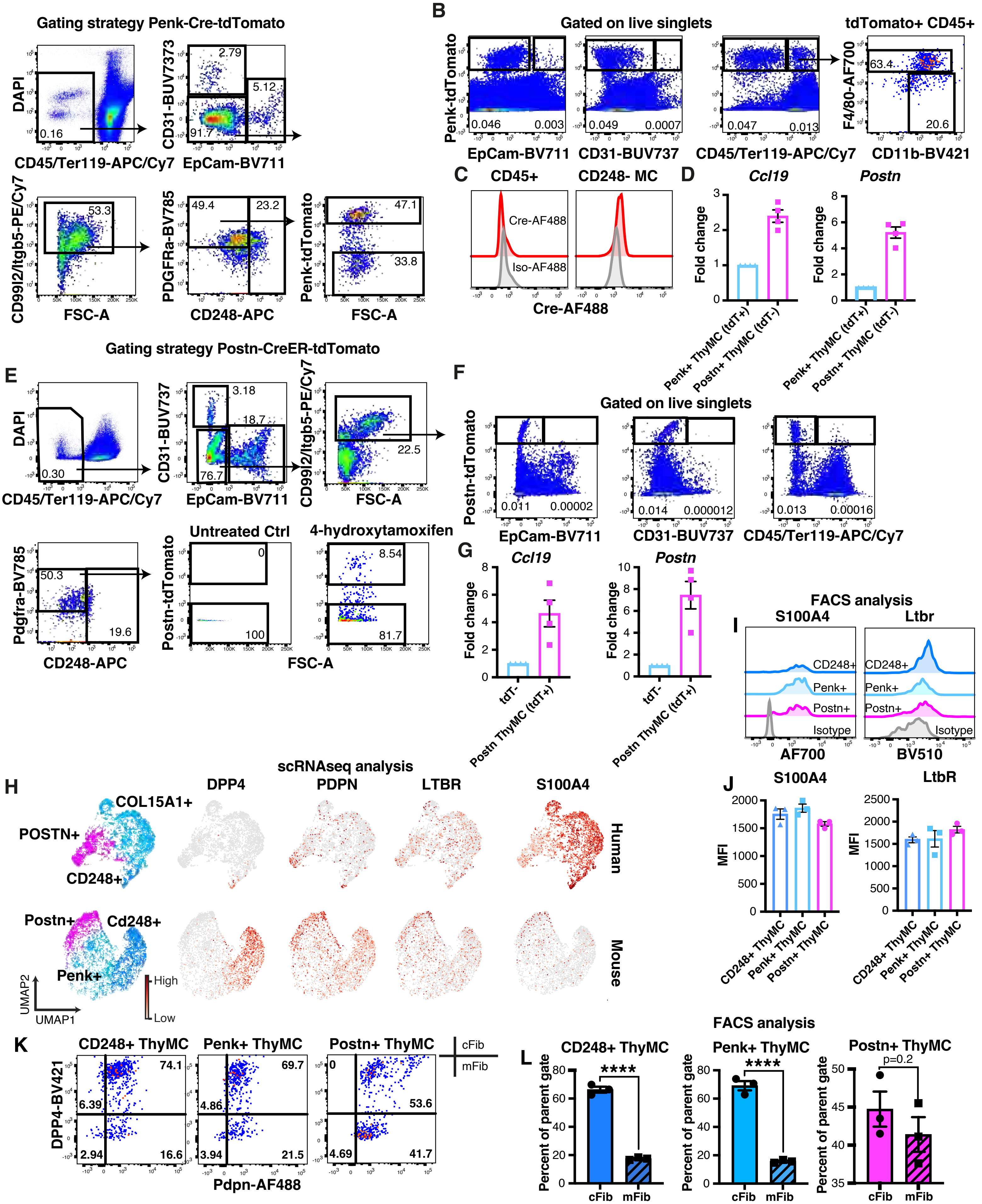

#### Supplemental Fig 6

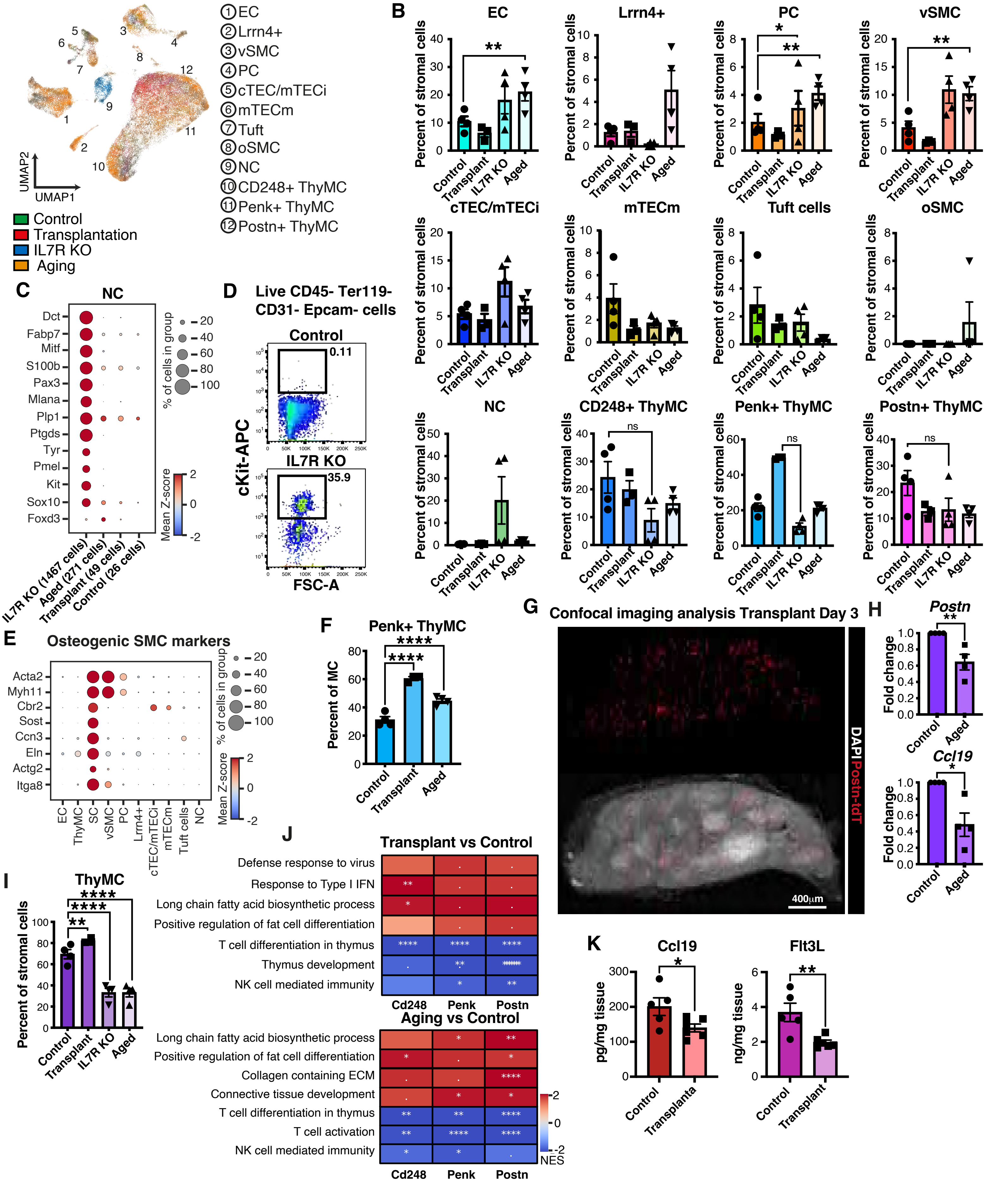

#### Supplemental Fig 7

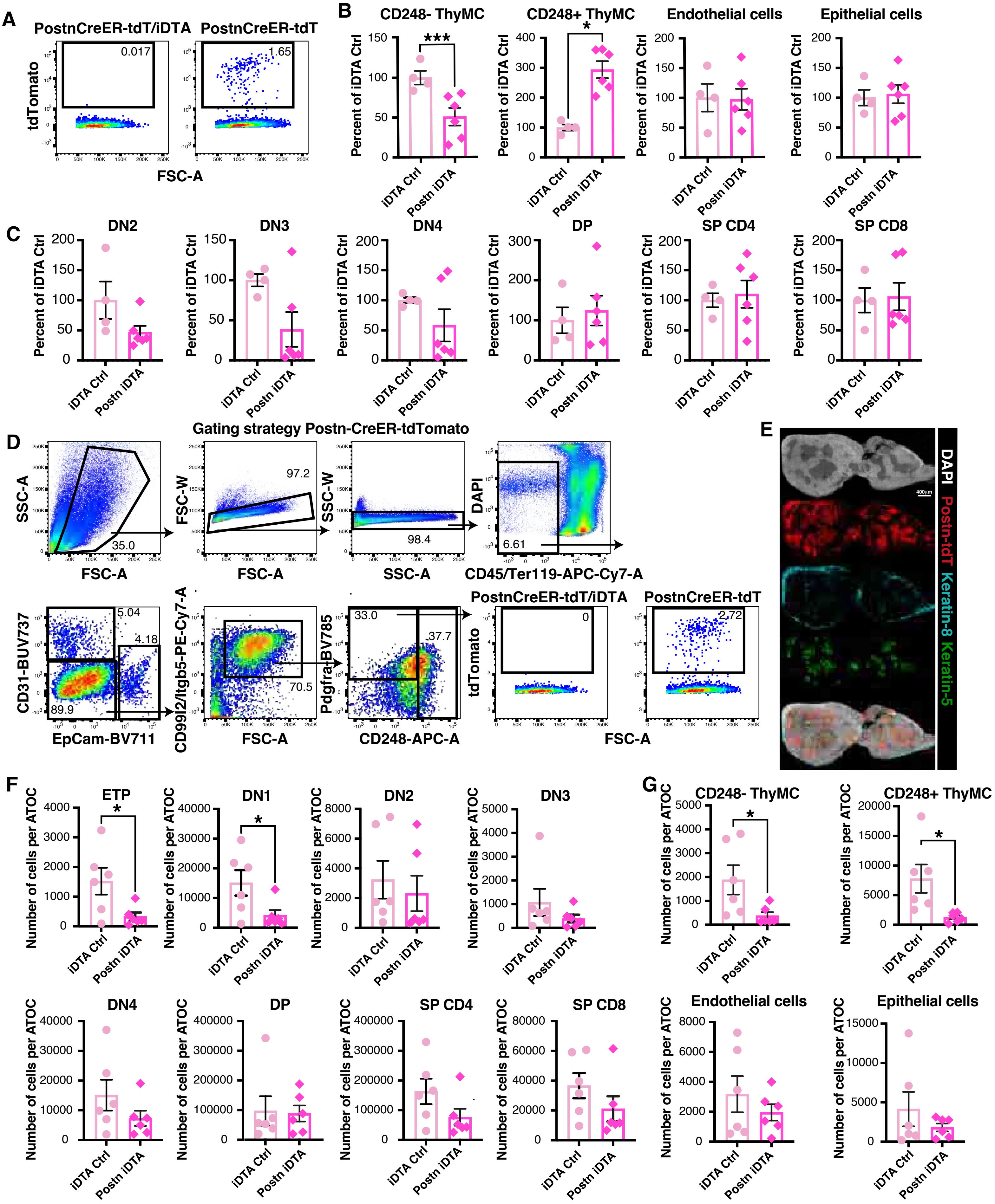

### Supplemental Fig 8

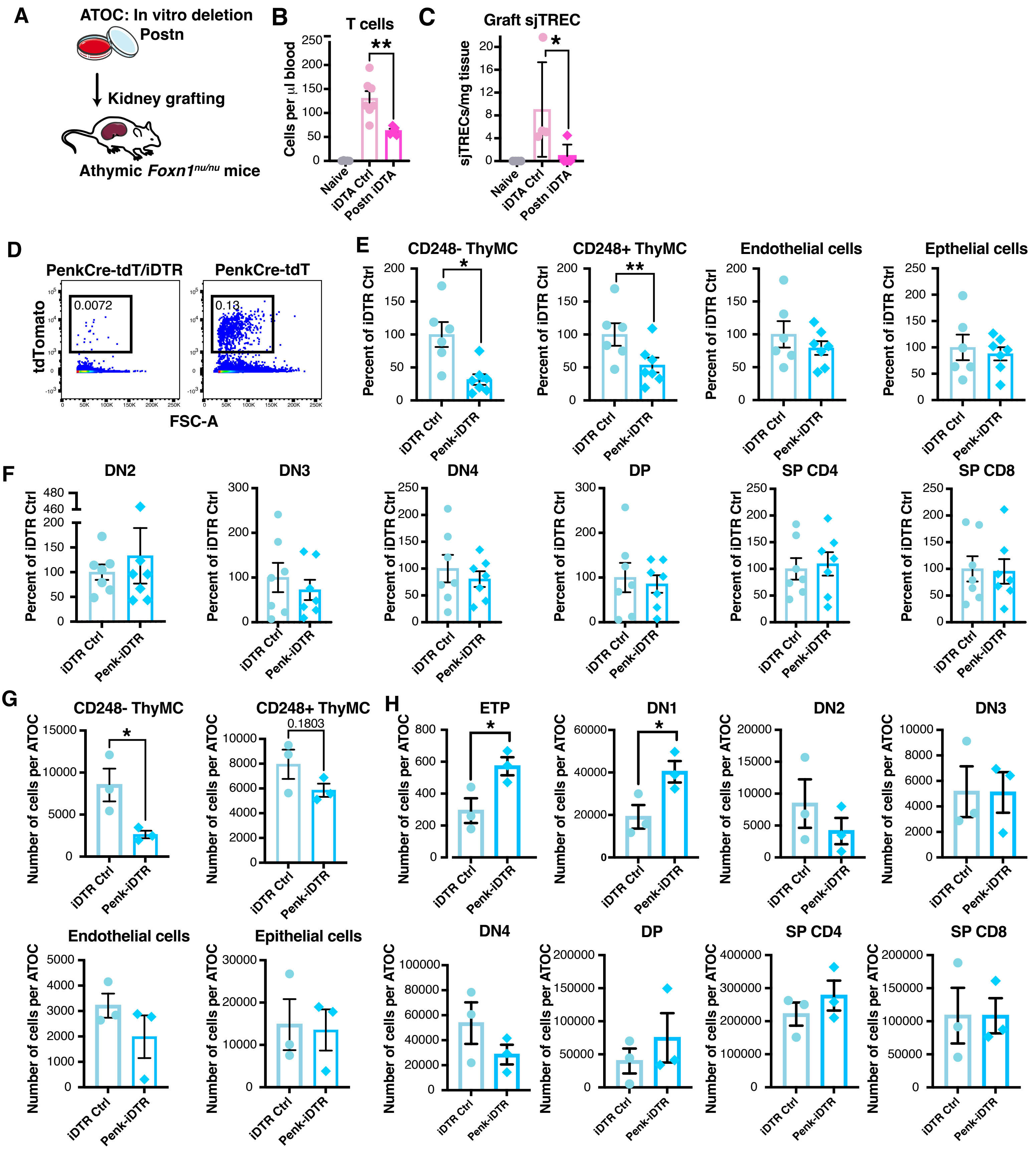

### Supplemental Fig 9

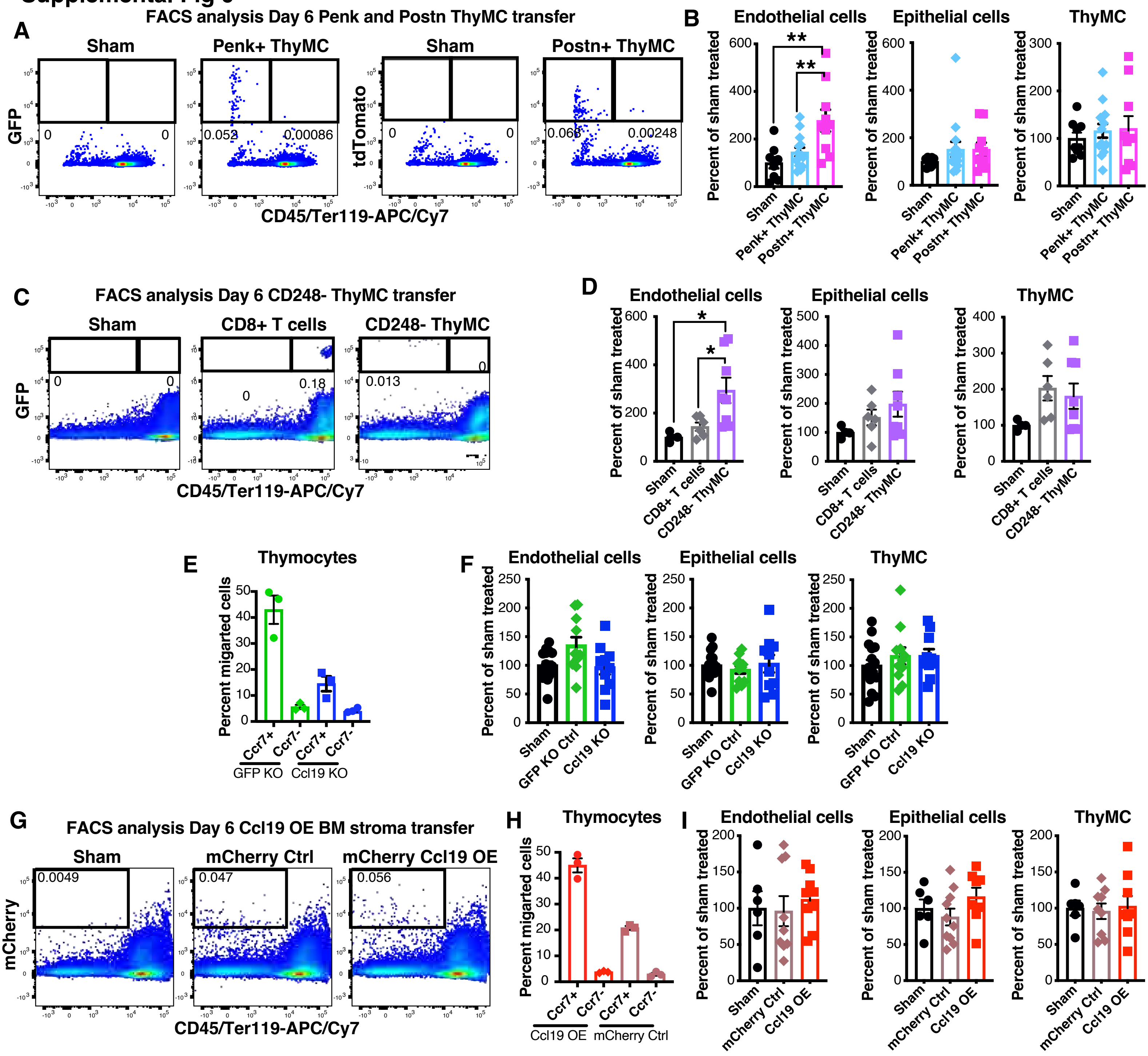

#### Supplemental Fig 10

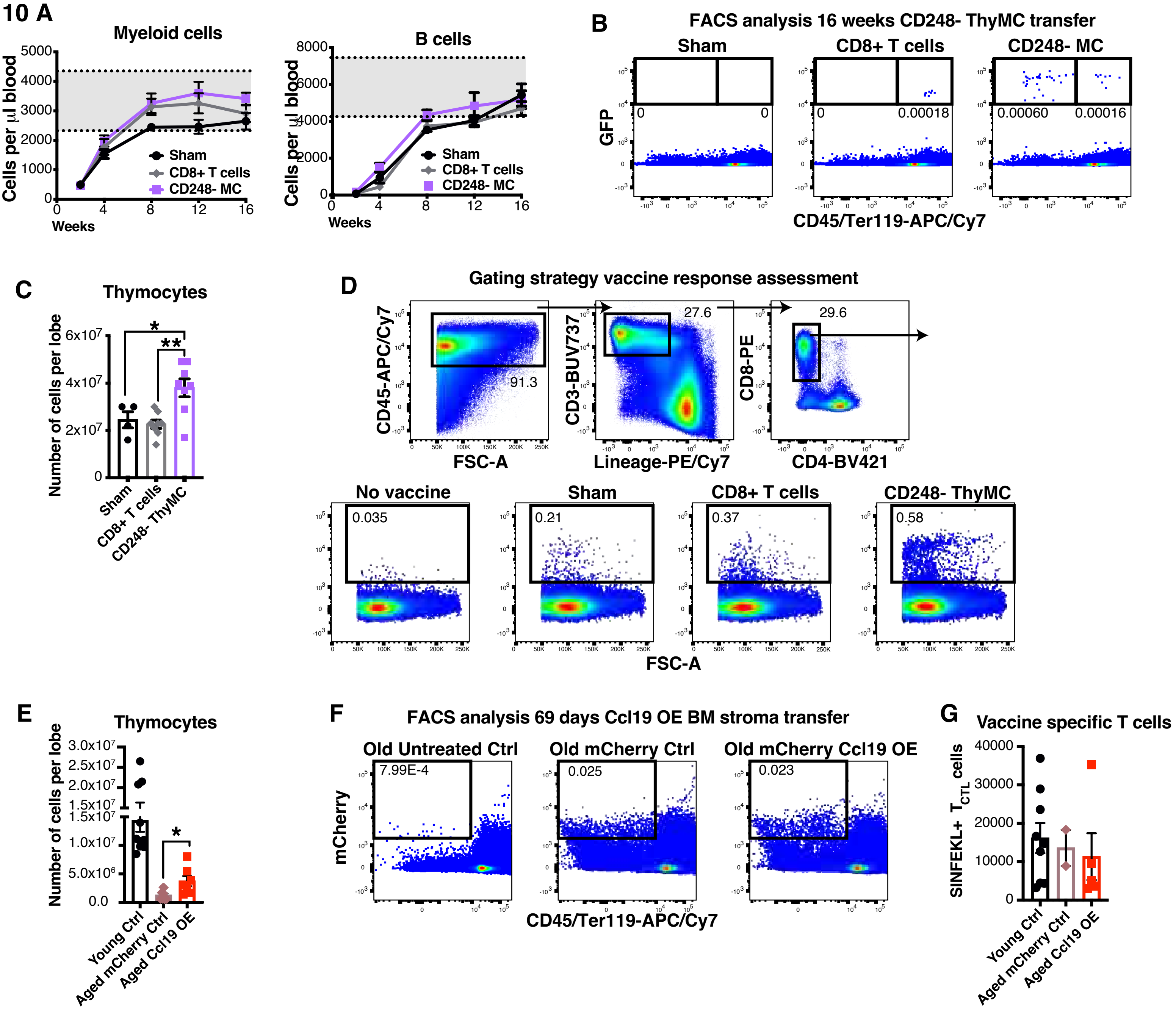
